# Targeting pathogenic VWF/ADAMTS13 dysregulation attenuates CTEPH progression

**DOI:** 10.64898/2026.06.17.732997

**Authors:** Zhijian Wu, Huan Dong, Quan Zhang, Zhirong Chai, Anlun Li, Esteban M. Dominguez, Xinyang Zhao, Leslie Spikes, Michael J Soares, X. Long Zheng, Liang Zheng

**Author notes:** Correspondence should be sent to: X. Long Zheng, Department of Pathology and Laboratory Medicine, The University of Kansas Medical Center, 3901 Rainbow Blvd, Delp 5016, Kansas City, KS 66160, USA, or Liang Zheng, Department of Pathology and Laboratory Medicine, The University of Kansas Medical Center, 3901 Rainbow Blvd, MS 3050, Kansas City, KS 66160, USA.

## Abstract

Chronic thromboembolic pulmonary hypertension (CTEPH) is a life-threatening pulmonary vascular disease, characterized by persistent thrombotic obstruction and progressive pulmonary vascular remodeling, yet the molecular mechanisms linking persistent thrombosis to vascular remodeling remain incompletely understood. Clinical studies have reported elevated plasma von Willebrand factor (VWF) levels and reduced ADAMTS13 in patients with CTEPH, but whether VWF/ADAMTS13 dysregulation contributes directly to disease pathogenesis remains unclear. Here, using newly established rat models of CTEPH, we identify a causative role for dysregulation of the VWF–ADAMTS13 axis in chronic thromboembolic progression. CTEPH rats developed persistent, unresolved VWF- and fibrin-rich thrombi accompanied by markedly increased endothelial VWF deposition. In contrast, ADAMTS13 expression and activity were significantly reduced in CTEPH rats. Consistent with these findings, genetic *Adamts13* deficiency further exacerbates pulmonary microvascular thrombosis and accelerated early mortality following disease induction. Mechanistically, ultra-large (UL)-VWF accumulated on the pulmonary endothelial surface, promoting robust platelet recruitment under shear. This platelet–VWF interaction stimulated the release of platelet-derived pro-remodeling mediators, including TGF-β1 and PDGF-BB. Genetic ablation of *Vwf* markedly reduced in situ microvascular thrombosis within pulmonary arterioles, attenuated pulmonary arterial remodeling, and improved pulmonary hemodynamics. Moreover, treatment with recombinant ADAMTS13 reduced endothelial UL-VWF accumulation, suppressed platelet activation, and effectively prevented thrombosis and platelet-driven pro-remodeling signaling in CTEPH rats. Collectively, these findings identify dysregulation of the VWF–ADAMTS13 axis as a key driver of pulmonary thrombosis and vascular remodeling in CTEPH and support therapeutic targeting of this pathway as a potential disease-modifying strategy.

**Key Points:**

1. VWF-ADAMTS13 dysregulation promotes persistent pulmonary thrombosis and platelet-driven arterial remodeling within pulmonary arterioles.
2. Recombinant ADAMTS13 treatment or VWF ablation abrogates pulmonary arterial thrombosis and halts vascular remodeling in CTEPH.

## Introduction

Pulmonary hypertension (PH) affects approximately 1% of the population globally and up to 10% of individuals aged >65 years(1). It is characterized by endothelial dysfunction, excessive vasoconstriction, pulmonary arterial remodeling, and in situ thrombosis, leading to increased pulmonary arterial pressure and resistance, progressive right ventricular failure and premature death(2),(3, 4). Chronic thromboembolic pulmonary hypertension (CTEPH), a severe subtype of PH, arises from unresolved pulmonary emboli(5–7). Despite adequate anticoagulation, approximately 5–10% of patients with acute pulmonary embolism (PE) progress to CTEPH(8–10). This underscores the necessity and importance to better understand the mechanisms underlying this disease progression and identify new therapeutic targets.

To date, limited treatment options are available for CTEPH. The most effective treatment for CTEPH is pulmonary thromboendarterectomy, a surgical procedure that removes organized thrombi from the pulmonary arteries(11–13). However, a substantial proportion of patients are not suitable candidates for surgery because of distal thrombus location and/or comorbid conditions(14). For these individuals, alternative treatments include balloon pulmonary angioplasty, a minimally invasive procedure that uses balloon catheters to dilate the narrowed pulmonary arteries to improve blood flow and reduce pulmonary artery pressure(15, 16). Pharmacological treatments including soluble guanylate cyclase stimulator riociguat that is approved for managing inoperable or residual CTEPH to reduce symptoms and improve hemodynamics(17, 18). Life-long anticoagulation remains the fundamental component of therapy to prevent recurrent thromboembolic events(19). Thus, there is an urgent need to better understand the pathogenesis of PE and CTEPH and to develop more effective therapeutics to dissolve and prevent chronic thrombi, as well as improved strategies for disease stratification and intervention to improve patient outcomes.

In this study, we explore the critical role of von Willebrand factor (VWF), a large multimeric glycoprotein primarily synthesized by endothelial cells, and its key regulator ADAMTS13 (A disintegrin and metalloproteinase with thrombospondin type 1 repeats, member 13), in pathogenesis and therapeutic targeting of CTEPH. VWF is an important adhesion molecule that mediates platelet adhesion and aggregation at vascular injury sites via interaction with subendothelial collagen and platelet glycoprotein Ib (GPIb) receptor(20). Elevated plasma VWF levels have been reported in patients with CTEPH(21–23). ADAMTS13, a metalloproteinase primarily synthesized by hepatic stellate cells and released into the circulation, regulates VWF multimer size and activity through proteolytic cleavage of ultra-large VWF multimers (24, 25). Notably, plasma ADAMTS13 levels are significantly reduced in CTEPH patients (21). While clinical studies suggest the role of dysregulation of VWF- ADAMTS13 axis in pathogenesis of CTEPH, its causative role in disease process remains unknown. Our findings address this knowledge gap in the pathobiology of CTEPH and suggest that therapeutic strategies aimed at restoring the disrupted ADAMTS13–VWF balance may mitigate persistent pulmonary thromboembolism, attenuate progressive vascular remodeling, and ultimately reduce the risk of right heart failure and mortality.

## Methods

### Human plasma samples

EDTA-anticoagulated plasma samples were from patients with idiopathic pulmonary arterial hypertension (IPAH) and CTEPH. Age- and sex-matched healthy control plasma samples were also collected from volunteer donors. All procedures involving human specimens were approved by the Institutional Review Board of the University of Kansas Medical Center (STUDY00145817).

### Animals

All animal experimental procedures were approved by the institutional animal care and use committee at the University of Kansas Medical Center (Protocol #2022-08-259). *Adamts13^-/-^* rats were established in our laboratory using CRISPR/Cas9 and fully characterized as described elsewhere(26). The *Vwf-*deficient rats provided by the laboratory of Dr. Robert R. Montgomery at the Versiti Blood Research Institute(27).

### Establishment of a rat model of CTEPH

CTEPH in rats was established by a repeated intrajugular veinous administration of autologous clots (Supplemental methods) once a week for 4 weeks with a subcutaneous injection of SU5416 (20 mg/kg), a vascular endothelial growth factor receptor 2 (VEGF2) inhibitor in the first week.

### Hydrodynamic measurement of pulmonary arterial pressure

The right ventricular systolic pressure (RVSP) was determined as the terminal procedure one week after the last clot administration via a catheter following anesthesia.

### Echocardiographic measurement

Transthoracic echocardiography was performed in rats using a high-resolution ultrasound system (Vevo F2, VisualSonics) equipped with a linear-array transducer.

### Microfluidic shear-based assay

Microfluidic channels (Fluxion Bioscience) were coated with type I fibrillar collagen (100 μg/mL) and blocked with 0.5% bovine serum albumin (BSA). Rat whole blood anticoagulated with PPACK (100 μM) was labeled with fluorescent anti-CD61 antibody and perfused over collagen-coated channels at arterial shear (50 dyne/cm²). Thrombus formation was recorded every 3 seconds for 180 seconds.

### Flow cytometry

Whole blood was collected into ACD and platelet-rich plasma was isolated by centrifugation (100 × g, 10 min). Platelets were pelleted (800 × g, 10 min), resuspended in calcium-free Tyrode’s buffer with prostaglandin E1 (1 μM), and stained with FITC–anti-CD61 and PE–anti-CD62P antibodies for 20 min at room temperature in the dark. Samples were analyzed immediately on a Cytek Aurora flow cytometer.

### Histological and immunohistochemical analysis

All histological analyses were performed in fixed and paraffin-embedded tissue sections. Immunohistochemical studies were conducted on both cryosections and paraffin-embedded tissue sections following manufacturers’ recommendations with some modifications.

### Single-cell RNA sequencing

Lung tissues were harvested and cells were dissociated and processed to the 10x Genomics Chromium platform workflow for single cell RNA sequencing.

### Soluble biomarker analysis

Human plasma ADAMTS13 antigen was measured by a commercial ELISA kit, and recombinant ADAMTS13 activity was assessed using the FRETS-VWF73assay(28). Rat plasma ADAMTS13 activity was determined by cattle VWF71 assay. Plasma NT-proBNP, D-dimer, TGF-β1 and PDGF-BB levels were quantified by ELISA according to the manufacturers’ protocols.

### In situ hybridization

In situ detection of *Adamts13* and *Vwf* mRNA expression was carried out with paraffin-embedded tissue sections using RNAscope probes specific for rat *Adamts13* or rat *Vwf* (ACD Bio, Newark, CA) and detected by fluorescent labeling technique.

### Statistical Analysis

Statistical analyses were performed using Prism 10 (Graphpad). Data are presented as mean ± SEM. Two-group comparisons used two-tailed Student’s t test for data that passed the normality test or the Mann-Whitney rank-sum U test for data that were not normally distributed. One-way analysis of variance (ANOVA) with post hoc Tukey’s test was used to evaluate differences among groups when three or more groups were analyzed. Two-way mixed-effects ANOVA followed by Tukey’s test was used for experiments involving repeated measurements and multiple groups. A P-value less than 0.05 was considered statistically significant.

*More detailed methods in Supplementary materials*.

## Results

### Elevated plasma VWF multimer levels in CTEPH patients

To analyze the structure and size distribution of VWF multimers in CTEPH, we performed VWF multimer pattern assays in plasma samples from healthy controls, patients with idiopathic pulmonary arterial hypertension (IPAH), and those with CTEPH (**Figure 1A**). Detailed demographic and clinical information for all participants is provided in **Supplementary Table S4**. The results revealed that the total intensity of plasma VWF (**Figure 1B**) and the ratio of high to low molecular weight VWF multimers (**Figure 1C**) were significantly increased in CTEPH patients compared with healthy controls, while no statistically significant difference was observed between IPAH patients and healthy controls, supporting the hypothesis that a dysregulation in the VWF-ADAMTS13 axis may contribute to the pathogenesis of CTEPH.

**Figure 1.**
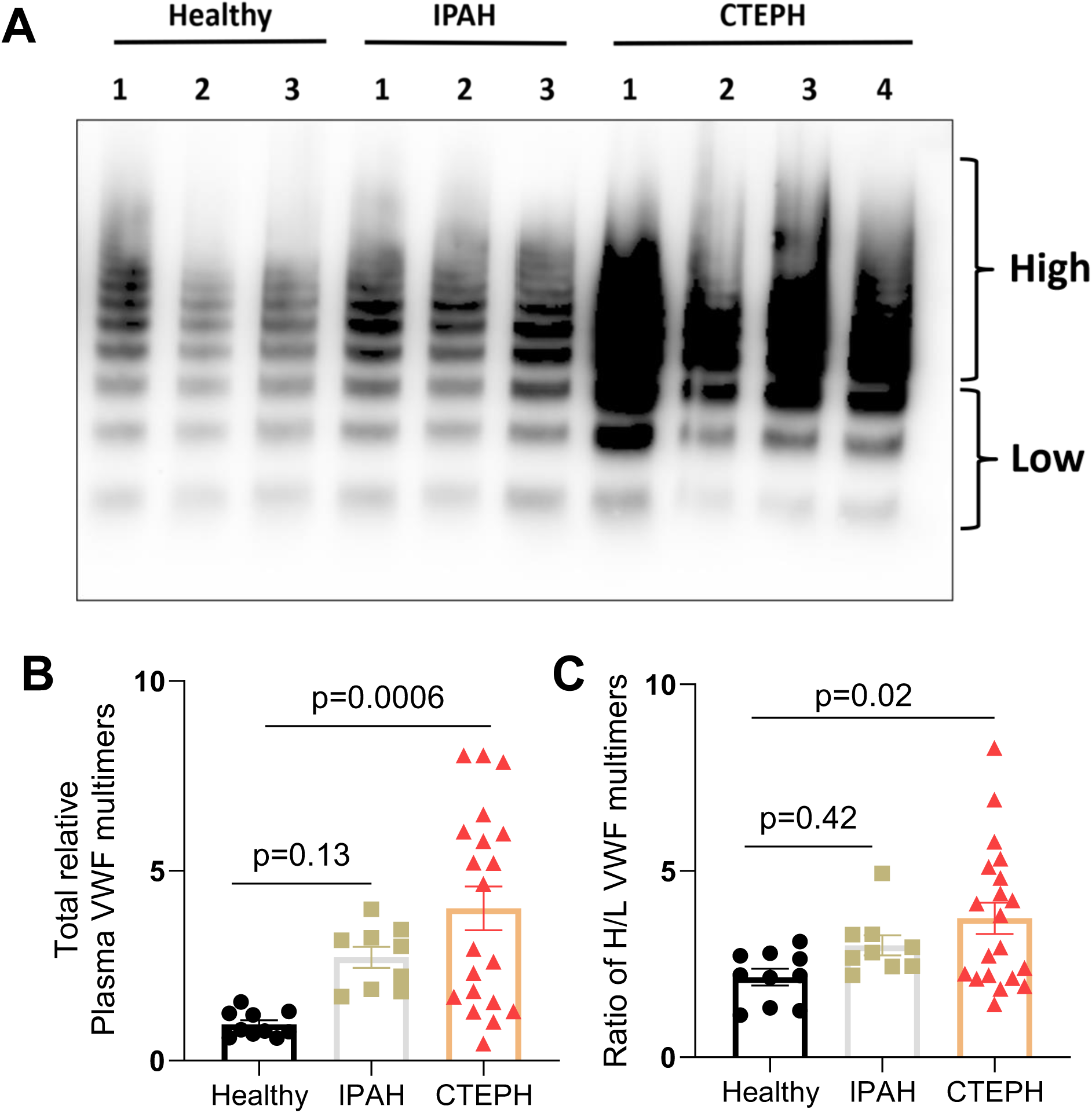
Elevated plasma levels of ultra-large VWF multimers in CTEPH patients. **A, B,** and **C** show plasma VWF multimer distribution, total multimer density, and the ratios of high to low VWF multimers, respectively, in patients with CTEPH compared to healthy controls and patients with idiopathic pulmonary hypertension (IPAH). Data shown are the individuals (dots), means (bars) ± standard errors of the means (SEM) (horizontal lines). The statistical significance of the differences among three groups was determined by one-way ANOVA followed by the Tukey’s test. P values less than 0.05 and 0.01 are considered statistically significant and highly significant, respectively.

### Rat model of CTEPH recapitulates the key hemodynamic and pathological features of human CTEPH

To establish a robust rat model of CTEPH, we systematically compared different modeling strategies (**Figure 2A).** Five different experimental groups were included. Sham: jugular vein injection of saline; SU: subcutaneous SU5416 injection; Clots (w/SU): jugular vein injection of autologous clots pre-conditioned with SU5416; Clots+SU: autologous clot injection combined with subcutaneous SU5416; and Clots(w/SU)+SU: injection of SU5416-pre-conditioned autologous clots combined with subcutaneous SU5416. Autologous clots were injected weekly via a jugular vein for four consecutive weeks, while subcutaneous SU5416 was delivered once on day 1. As shown in **Figure 2B-C**, after 4 weeks experimental induction, both the Clots + SU5416 group and the Clots (w/SU) + SU5416 group developed dramatically elevated right ventricular systolic pressure (RVSP) and significant right ventricular hypertrophy (Fulton index), whereas the rats receiving SU5416 alone or SU5416-containing autologous clots alone exhibited only modest or no increases in RVSP (**Figure S1A**). Therefore, the Clots + SU5416 model, which employs autologous clots without prior SU5416 preconditioning to induce CTEPH, better mimics the spontaneous development of obstructive thrombi and was therefore selected for subsequent studies.

**Figure 2.**
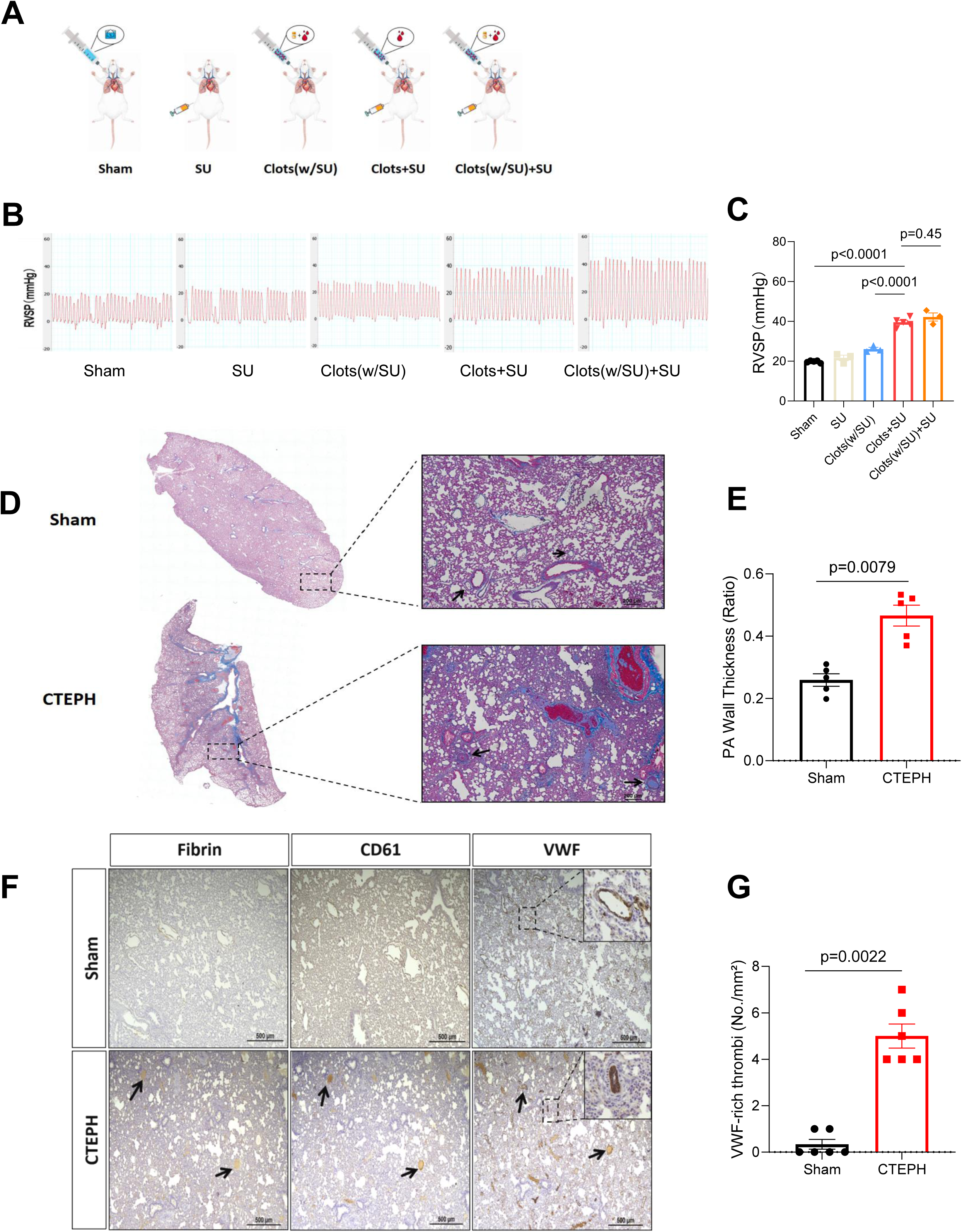
Rat model of CTEPH recapitulates the key hemodynamic and pathological features of human CTEPH. **A.** Schematic representation of the experimental strategies. Sham: jugular vein injection of saline; SU: subcutaneous SU5416 injection; Clots (w/SU): jugular vein injection of autologous clots pre-conditioned with SU5416; Clots+SU: autologous clot injection combined with subcutaneous SU5416; and Clots(w/SU)+SU: injection of SU5416-pre-conditioned autologous clots combined with subcutaneous SU5416. Autologous clots were injected weekly via a jugular vein for four consecutive weeks. Subcutaneous SU5416 was delivered once on day 1. Blood was collected at baseline for clot formation, and terminal blood collection, hemodynamic measurements, and tissue harvest were performed on day 28. **B** and **C** show the tracing and quantification, respectively, of the right ventricular systolic pressure (RVSP) measured through right heart catheterization in anaesthetized rats of various groups. **D** and **E** show the Masson’s trichrome staining of lung sections and pulmonary arteriole wall thickness, respectively, from Sham and CTEPH rats (Clots + SU). **F**. Immunohistochemical staining of lung sections from CTEPH and Sham rats for fibrin, platelet CD61, and VWF. **G.** Quantification of occlusive VWF-rich pulmonary thrombi per mm² in Sham and CTEPH lungs. Data shown are individual values (dots), means (bars) ± standard errors of the means (SEM) (horizontal lines). The statistical analysis for the differences among three groups or more was performed with one-way ANOVA followed by the Tukey’s test or two-tailed unpaired t-test was perform comparing the difference in two groups. P values less than 0.05 and 0.01 are considered to be statistically significant and highly significant, respectively.

Histological analysis of lungs via Masson’s trichrome staining revealed the presence of organized thrombi within pulmonary arteries, accompanied by luminal narrowing and increased vessel wall thickness in Clots + SU5416 rats, the changes that were absent in Sham controls (**Figure 2D**-**E****, Figure S1B**). Echocardiography further validated this CTEPH phenotype through impaired pulmonary hemodynamics, as reflected by the reduced PAT/PET ratio (pulmonary acceleration time to pulmonary ejection time ratio, a non-invasive echocardiographic marker used to evaluate pulmonary artery pressure and right ventricular function) and increased RVWTD (right ventricular wall thickness in diastole, a key parameter for identifying structural changes in the heart) (**Figure S1C–E**).

Immunohistochemical staining showed widespread and persistent microvascular thrombosis in CTEPH lungs, with pulmonary arterioles exhibiting strong positivity for fibrin, CD61, and VWF, indicative of platelet-rich and VWF-associated thrombi retained within the pulmonary circulation (**Figure 2F-G**). These findings were supported by markedly elevated plasma levels of D-dimer (**Figure S1F**) and NT-proBNP (**Figure S1G**) in CTEPH rats compared to the controls, reflecting ongoing thrombotic activity and significant myocardial stress. Together, these results indicate the successful establishment of a new rat CTEPH model that recapitulates the key hemodynamic and pathological features of human CTEPH, including pulmonary atrial hypertension and associated right ventricular remodeling, pulmonary vascular obstruction and microvascular thrombosis, as well as vascular remodeling.

### Dysregulation of VWF-ADAMTS13 axis in the CTEPH rats

Lung single-cell RNA sequencing demonstrated that *Vwf* transcription was selectively and robustly upregulated in pulmonary arterial endothelial cells (aECs), whereas no significant changes were observed in venous, lymphatic, or capillary endothelial subpopulations in CTEPH lungs (**Figure 3A-B**). To explore the functional implications of this upregulation, we evaluated localized VWF levels following disease induction. Immunohistochemical analysis confirmed a significant increase in VWF expression across both large and small pulmonary arteries in CTEPH rats, while pulmonary veins remained unaffected (**Figure 3C-D**). Furthermore, immunofluorescence staining revealed a marked accumulation of luminal VWF strings anchored to the pulmonary arterial wall in the CTEPH cohort compared to the Sham group (**Figure 3E-F**). These data indicate a substantial increase in arterial VWF expression coupled with compromised proteolytic cleavage of VWF on the endothelial surface during CTEPH pathogenesis. Consistent with human CTEPH data, CTEPH rats exhibited significantly elevated plasma VWF levels enriched with high-molecular-weight VWF multimers (**Figure 3G–I**). Notably, SU5416 alone did not alter VWF expression or multimer distribution (**Figure S2**). Western blot analysis of lung homogenates confirmed significantly increased overall VWF protein and marked P-Selectin upregulation in CTEPH rats compared with Sham controls, indicating enhanced endothelial activation (**Figure S3A-C**). Furthermore, VWF- and fibrin-rich thrombi were prevalent within pulmonary arterioles of CTEPH lungs (**Figure S3D**).

**Figure 3.**
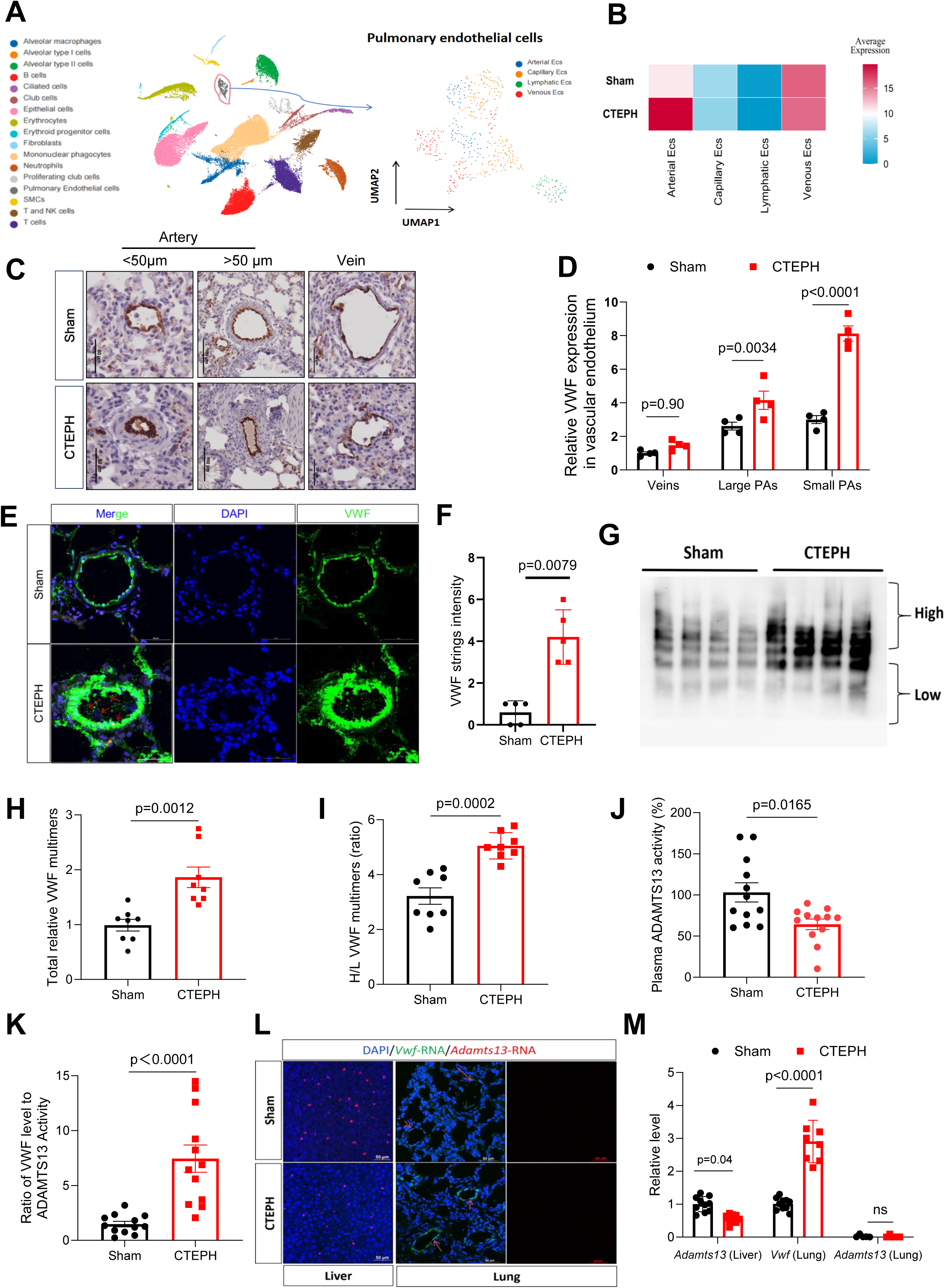
VWF-ADAMTS13 dysregulation in the rat model of CTEPH. **A**. Uniform manifold approximation and projection visualization of lung endothelial cells identified by single-cell RNA sequencing of dissociated left lungs from Sham and CTEPH rats Endothelial cells were further classified into arterial, capillary, venous, and lymphatic endothelial subtypes. **B**. Heatmap showing the average VWF transcript expression across various endothelial subtypes from Sham and CTEPH rats. **C**. Representative immunohistochemical staining of VWF in pulmonary veins and large (>50 μm) and small (<50 μm) pulmonary arteries from Sham and CTEPH rats. **D**. Quantitative analysis of relative VWF expression in pulmonary veins and large and small pulmonary arteries, in Sham and CTEPH rats. **E.** Immunofluorescence staining of pulmonary arteries from Sham and CTEPH rats showing VWF (green) and nuclei with DAPI (blue). UL-VWF strings were stuck along the luminal surface of pulmonary arteries in CTEPH rats (red arrows). **F**. Quantification of VWF string intensity in pulmonary arteries from Sham and CTEPH rats. **G, H,** and **I**. Representative plasma VWF multimer distribution patterns, relative VWF density, and the ratio of high to low molecular weight multimers, respectively, from Sham and CTEPH rats. **J**. Plasma ADAMTS13 activity measured prior to and following CTEPH induction. **K**. The ratios of plasma VWF to ADAMTS13 activity in Sham and CTEPH rats. **L**. Representative in situ hybridization images showing *Vwf* and *Adamts13* mRNA expression in lung tissue of Sham and CTEPH rats. **M**. Relative fluorescence signals for *Vwf* and *Adamts13* mRNA by in situ in lung and liver tissues of Sham and CTEPH rats. Data presented are individual values (dots), means (bars) ± standard errors of the means (SEM) (horizontal lines). An unpaired two-tailed Student’s t-test was performed to determine the difference between two groups. P values less than 0.05 and 0.01 are considered to be statistically significant and highly significant, respectively.

We next assessed whether this VWF accumulation resulted from diminished ADAMTS13. Plasma ADAMTS13 activity was significantly reduced in CTEPH rats, leading to a significantly increased ratio of VWF/ADAMTS13 activity (**Figure 3J-K**). *In situ* hybridization demonstrated that *Adamts13* mRNA is predominantly expressed in the liver, with minimal to undetectable expression in the lung tissues (**Figure 3L**). Lung single-cell RNA sequencing analysis also confirmed that *Adamts13* expression was not derived from pulmonary endothelial cells (**Figure S2E**). Instead, hepatic *Adamts13* mRNA expression was significantly reduced in CTEPH rats compared with sham controls, while lung *vwf* mRNA expression was markedly increased in CTEPH rats (**Figure 3M**, **Figure S2F-G**). Collectively, these findings indicate that excessive pulmonary arterial endothelial VWF production coupled with reduced hepatic ADAMTS13 expression dysregulates VWF-ADAMTS13 axis and impaired UL-VWF cleavage, likely leading to persistent pulmonary thrombosis, vascular remodeling in CTEPH rats.

### Adamts13 deficiency exacerbates pulmonary microvascular thrombosis and early death following CTEPH induction

To investigate if ADAMTS13 deficiency would directly affect the disease severity, we generate an *Adamts13* deficient rat using CRISPR/Cas9. Since ADAMTS13 is primarily secreted by hepatic stellate cells, this *Adamts13* deficient model allow us to evaluate the total systemic contribution of ADAMTS13 to the pulmonary thrombo-inflammatory environment. We subjected WT (*Adamts13^+/+^) and Adamts13^-/-^* rats which generated from *Adamts13^+/-^* breeding to Sham or CTEPH procedures (**Figure 4A**). Successful knockout of *Adamts13* was confirmed by genotyping and loss of hepatic ADAMTS13 protein expression (**Figure S4A-B**). Circulating ADAMTS13 antigen and activity were also undetectable in the Adamts13 knockout rats(26). Under basal conditions, pulmonary VWF protein expression remained unchanged in *Adamts13^-/-^* rats, indicating that *Adamts13* deficiency itself does not affect VWF synthesis in the lung (**Figure S4C-D**). Following CTEPH induction, both *Adamts13^+/+^ and Adamts13^-/-^* rats exhibited similar increases in RVSP and right ventricular hypertrophy, indicating a comparable hemodynamic burden (**Figure 4B–D**). Plasma VWF multimers were increased after CTEPH induction in both genotypes, with no significant difference in the proportion of high–molecular-weight VWF multimers between *Adamts13^+/+^ and Adamts13^-/-^* rats (**Figure 4E–G**). Despite similar hemodynamic severity, *Adamts13^-/-^* rats exhibited significantly higher early mortality compared with *Adamts13^+/+^*rats following CTEPH induction (**Figure 4I**). This was associated with a marked increase in pulmonary in situ microvascular thrombosis, characterized by enhanced fibrin- and CD61-positive thrombi (**Figure 4H** and **4J–K**). Pulmonary arterial wall thickness and arterial VWF levels (**Figure 4H, L–M**), plasma NT-proBNP and D-dimer levels did not differ significantly between *Adamts13^+/+^*and *Adamts13^-/-^* CTEPH rats (**Figure S4F–G**). These findings indicate that while ADAMTS13 deficiency does not further aggravate hemodynamics, it significantly increases local microvascular thrombosis and early mortality in CTEPH.

**Figure 4.**
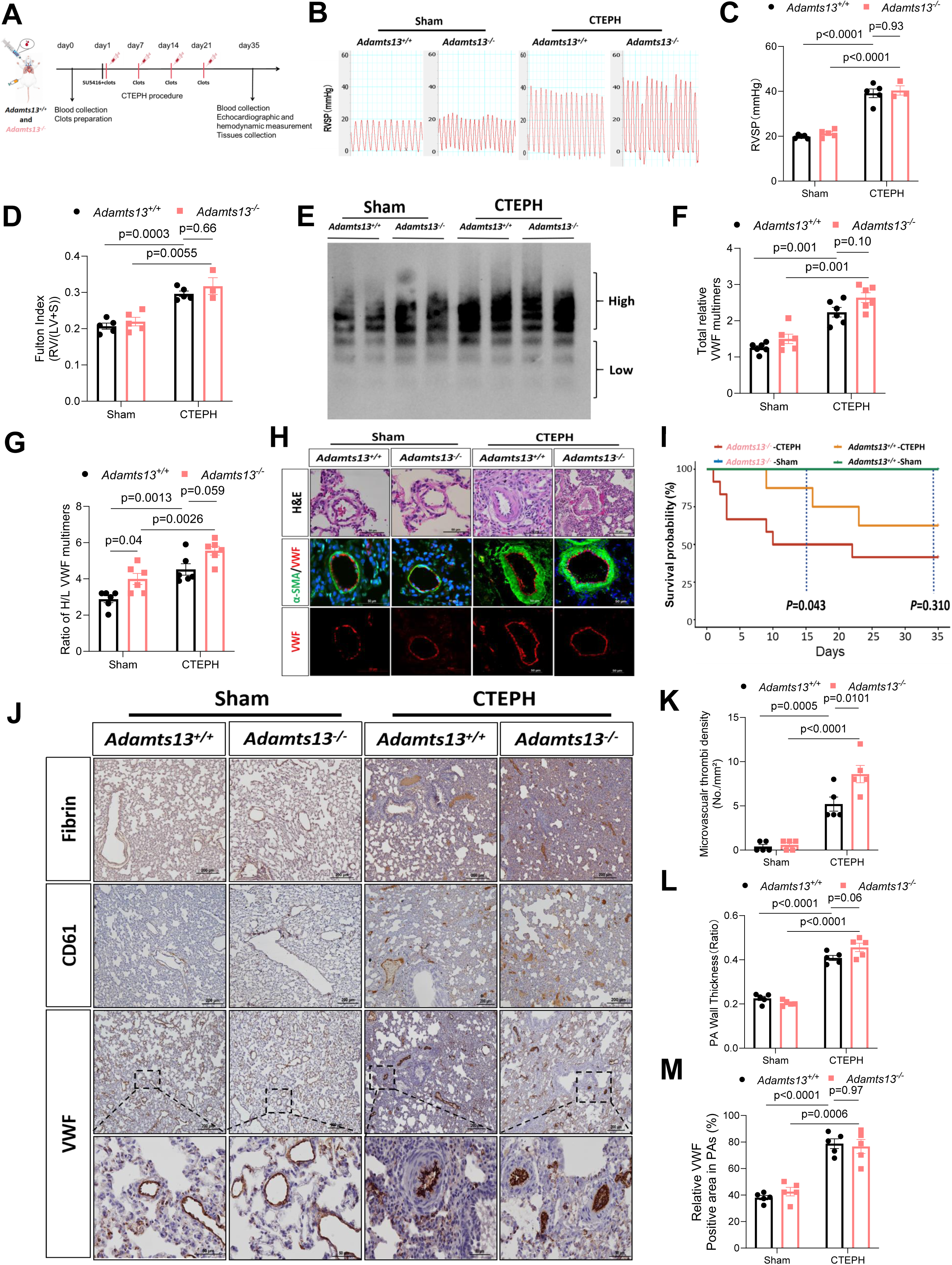
ADAMTS13 deficiency exacerbates pulmonary in situ microvascular thrombosis and results in early death following CTEPH induction. **A.** Two groups of rats (*Adamts13^+/+^ Adamts13^-/-^*) were subjected to Sham vs. CTEPH procedure. CTEPH was induced via repeated embolic challenges (weekly jugular vein delivery of autologous thrombi for 4 weeks) combined with systemic VEGFR2 inhibition (a single subcutaneous injection of SU5416 on Day 1). **B** and **C.** Right ventricular systolic pressure (RVSP) waveforms and quantitative analysis, respectively, in Sham and CTEPH rats with two different genotypes. **D.** Fulton index [RV/(LV+S)] in Sham and CTEPH rats. **E, F,** and **G.** Plasma VWF multimer distribution, total multimer intensity, and the ratio of high to low molecular weight multimers, respectively, in Sham vs. CTEPH rats. **H.** Representative pulmonary arterial H&E staining (top row) and immunofluorescence staining for α-SMA and VWF (middle row) or VWF alone (bottom row) in *Adamts13^+/+^ Adamts13^-/-^* rats with Sham vs. CTEPH procedure. **I.** Kaplan–Meier survival analysis of *Adamts13^+/+^* and *Adamts13^-/-^*rats with Sham vs. CTEPH procedure. **J.** Representative immunohistochemistry of lung tissue for fibrin, CD61, and VWF in *Adamts13^+/+^ Adamts13^-/-^* rats with Sham vs. CTEPH procedure. **K, L,** and **M.** Quantification of pulmonary microthrombi density, pulmonary arterial wall thickness ratio, and pulmonary arterial VWF intensity, respectively. Data shown are individual values (dots), means (bars), and standard errors of the means (SEM). The differences were analyzed by two-way ANOVA followed by the Tukey’s multiple comparisons test. P values less than 0.05 and 0.01 are statistically significant and highly significant, respectively.

### VWF deficiency attenuates pulmonary hypertension, thrombus formation and vascular remodeling in CTEPH rats

VWF is a circulating protein predominantly derived from endothelial cells. Given that both endothelial and circulating VWF levels were markedly elevated in CTEPH, we next investigated if VWF ablation would attenuate CTEPH phenotype. We subjected *Vwf^+/+^*, *Vwf^+/-^*and *Vwf*⁻*^/^*⁻ to Sham or CTEPH procedures (**Figure 5A**). Stepwise reductions in plasma VWF abundance and multimers were confirmed in *Vwf^+/-^*and *Vwf^-/-^* rats (**Figures S5A-B**). Following induction, *Vwf^-/-^* rats exhibited improved survival (**Figure S5C**) and a striking attenuation of pulmonary hypertension, evidenced by significantly reduced RVSP and improved PAT/PET ratios compared to *Vwf^+/+^*littermates (**Figure 5B-C, Figure S5D-E**). Right ventricular hypertrophy, assessed by both the Fulton index and RVWTD, was also markedly reduced in *Vwf*⁻*^/^*⁻ rats following CTEPH induction (**Figures 5D and S5F**). These structural and hemodynamic improvements were accompanied by significantly deceased plasma levels of NT-proBNP in *Vwf*⁻*^/^*⁻ rats compared with those in *Vwf^+/+^* controls after CTEPH induction, reflecting attenuated right ventricular strain by VWF deficiency (**Figure S5G**). Shear-based microfluidic assays demonstrated substantially impaired thrombus formation in blood from *Vwf*⁻*^/^*⁻ rats under both sham and CTEPH induction conditions (**Figure 5E and Figure S5I**). Notably, *Vwf^+/-^* rats also exhibited significantly delayed thrombus onset and reduced platelet adhesion and aggregation compared with *Vwf^+/+^*groups after CTEPH induction (**Figure 5E and F**). Moreover, *Vwf*⁻*^/^*⁻ rats after CTEPH induction exhibited markedly reduced fibrin deposition and platelet accumulation within pulmonary vessels compared with *Vwf^+/+^*rats, as assessed by fibrin and CD61 immunostaining(**Figure 5J**). Quantitative analysis further revealed a significant reduction in pulmonary microvascular thrombi density in *Vwf*⁻*^/^*⁻ rats after CTEPH induction, with *Vwf^+/-^* rats exhibiting an intermediate protective phenotype (**Figure 5K**). In line with these findings, plasma levels of D-dimer were significantly reduced in both *Vwf*⁻*^/^*⁻ and *Vwf^+/-^*rats (**Figure S5H**). In addition to attenuated thrombosis, VWF deficiency also reduced pulmonary arterial remodeling as shown by significantly decreased arterial wall thickness in *Vwf*⁻*^/^*⁻ rats following CTEPH induction compared with *Vwf^+/+^* controls (**Figure 5L**). Altogether, these data clearly indicated that genetic ablation of *Vwf* attenuates CTEPH progression in rats.

**Figure 5.**
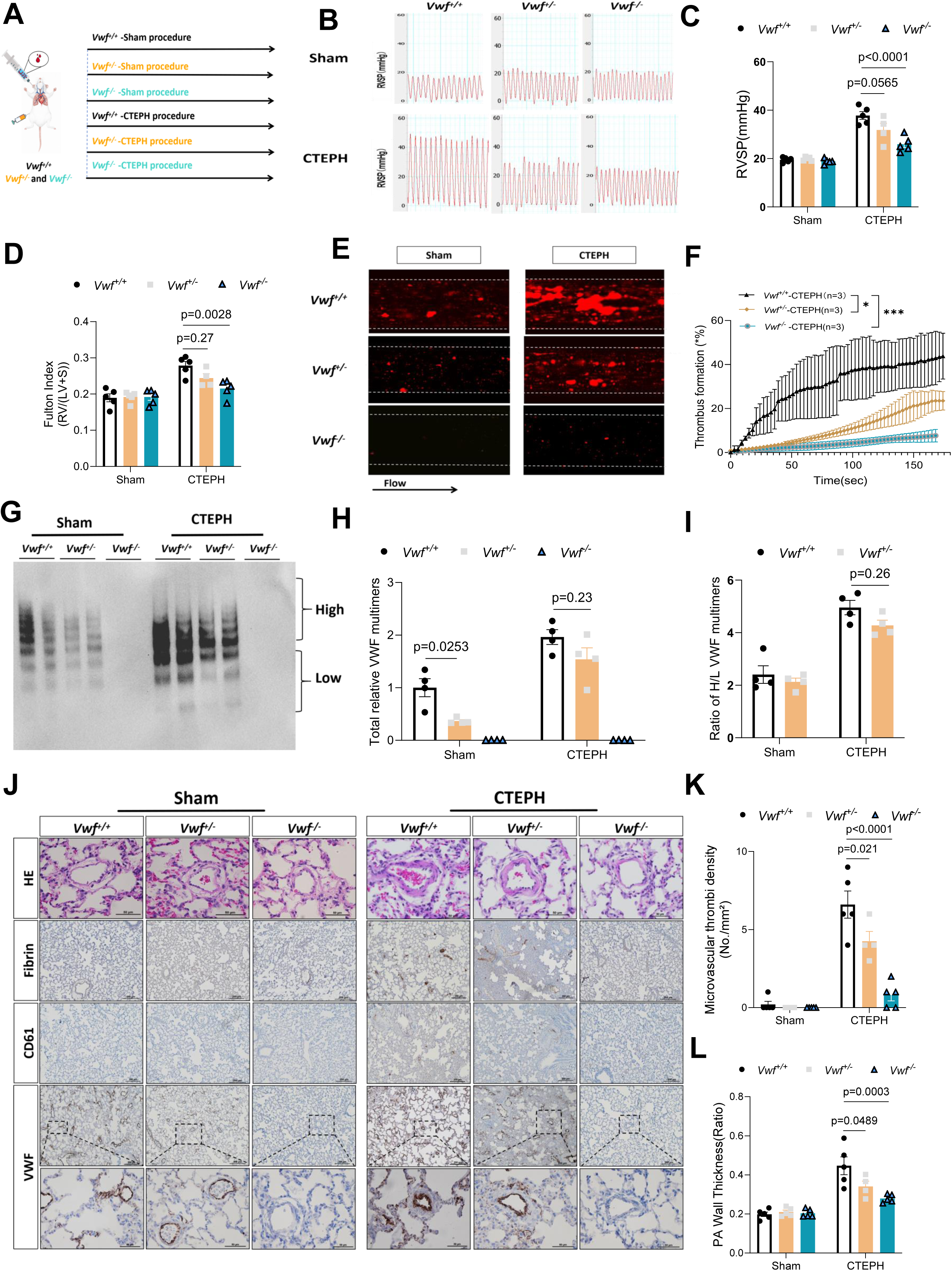
VWF deficiency attenuates pulmonary hypertension and thrombus formation in the rat model of CTEPH. **A.** Experimental design illustrating *Vwf^+/+^*, *Vwf^+/-^*and *Vwf*⁻*^/^*⁻ rats for Sham vs. CTEPH procedure. **B** and **C.** Representative right ventricular systolic pressure (RVSP) waveforms and quantitative analysis of RVSP, respectively, in rats with various genotypes (*Vwf^+/+^*, *Vwf^+/-^*and *Vwf*⁻*^/^*⁻) under Sham vs. CTEPH procedure. **D.** Fulton index [RV/(LV+S)] measured in *Vwf^+/+^*, *Vwf^+/-^* and *Vwf*⁻*^/^*⁻ rats following Sham vs. CTEPH procedure. **E** and **F.** Representative images and time-course quantitative analysis, respectively, of thrombus formation in a microfluidic based thrombosis assay using whole blood samples from *Vwf^+/+^*, *Vwf^+/-^* and *Vwf*⁻*^/^*⁻ rats under Sham vs. CTEPH procedure. **G, H,** and **I.** Plasma VWF multimer distribution, total plasma VWF multimer intensity, and the ratios of high to low molecular VWF multimers, respectively. **J.** Representative histological (top row) and immunohistochemical analyses or fibrin (2^nd^ row), CD61 (3^rd^ row), and VWF (bottom row) of lung sections from *Vwf^+/+^*, *Vwf^+/-^* and *Vwf*⁻*^/^*⁻ rats under Sham vs. CTEPH procedure. **K** and **L.** Quantitative analysis of pulmonary microthrombus density and pulmonary arterial wall thickness, respectively, in lung sections from *Vwf^+/+^*, *Vwf^+/-^*and *Vwf*⁻*^/^*⁻ rats under Sham vs. CTEPH. Data shown are individual values (dots), means (bars) ± standard erros of the means (SEM) (horizontal lines). the differences were analyzed by two-way ANOVA followed by the Tukey’s multiple comparisons test. For time-course thrombosis analysis, two-way repeated-measures ANOVA was applied. Here, * and *** indicate P values < 0.05 and < 0.005, respectively.

### VWF-mediated platelet adhesion promotes platelet activation and release of platelet-derived pro-remodeling factors under shear

To investigate the mechanistic link between UL-VWF, platelet activation, and pulmonary vascular remodeling, we examined platelet–VWF interactions using a microfluidic system. Under arterial flow, blood from *Vwf^+/+^*rats generated prominent thrombus-associated linear UL-VWF strings that supported platelet adhesion and robust P-selectin expression, which were largely absent in *Vwf*⁻*^/^*⁻ rat blood (**Figure 6A-B**). Supplementation of *Vwf*⁻*^/^*⁻ rat blood with purified human VWF restored UL-VWF string formation, platelet adhesion and activation, whereas treatment of *Vwf^+/+^* blood with a recombinant truncated ADAMTS13 efficiently disrupted UL-VWF strings and attenuated UL-VWF–dependent platelet recruitment and activation under shear (**Figure 6A-B, Figure S5J**).

**Figure 6.**
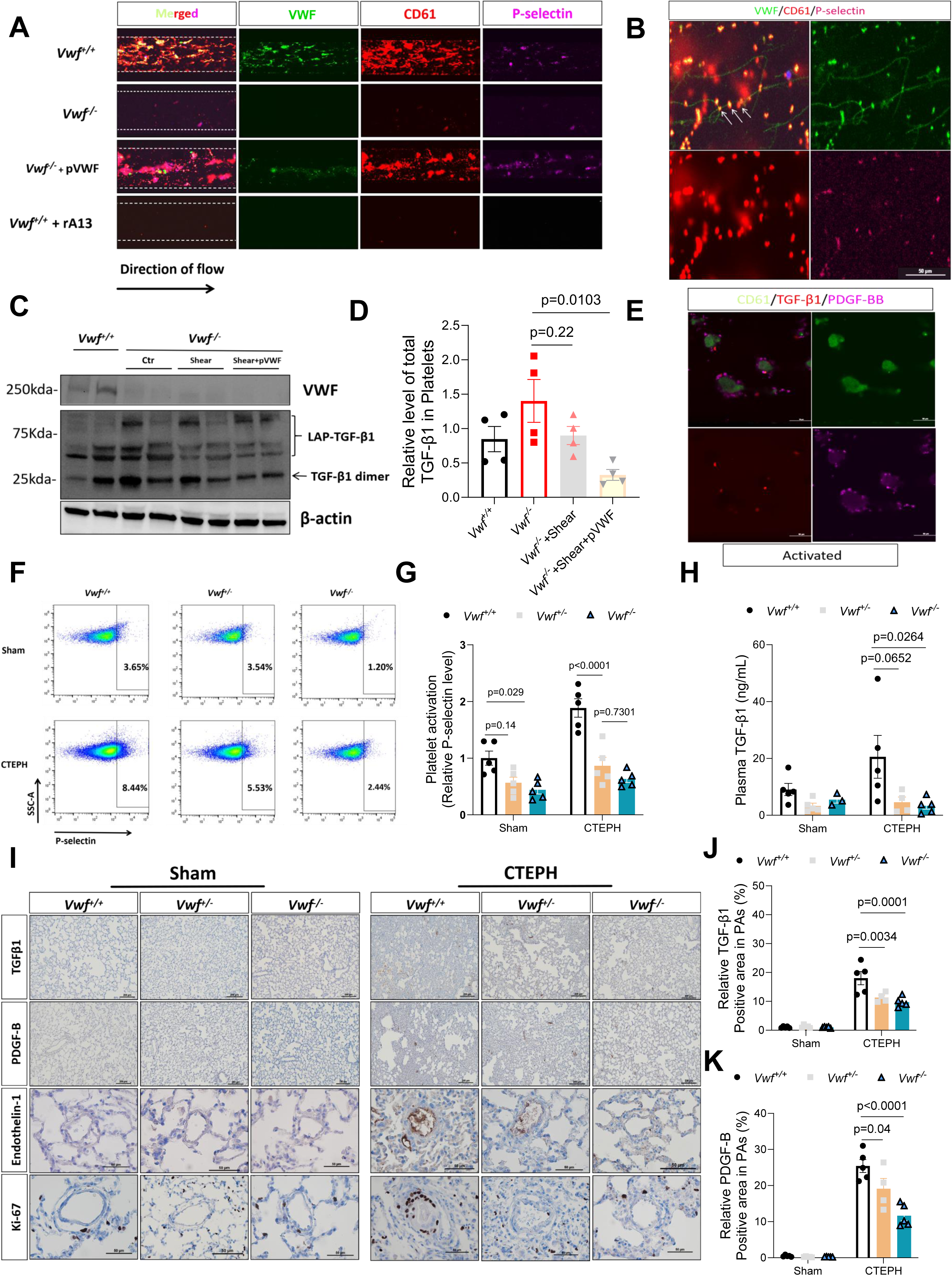
VWF-dependent platelet adhesion promotes platelet activation and the release of platelet-derived factors under shear. **A.** Representative immunofluorescence images from the microfluidic shear-based assay showing VWF (green), platelet marker CD61 (red), and P-selectin (magenta) using whole blood from *Vwf^+/+^*and *Vwf*⁻*^/^*⁻ rats, as well as *Vwf*⁻*^/^*⁻ blood supplemented with purified human VWF (100 µg/ml) or *Vwf^+/+^* blood treated with recombinant ADAMTS13 truncated after the 5^th^ TSP1 repeat (rA13 in short, activity: 0.32U/mL). Merged images are shown in the first column and the direction of flow is indicated with an arrow. **B.** Higher-magnification images illustrating the colocalization patterns of VWF, CD61, and P-selectin in thrombus-associated structures formed under shear conditions. Arrows indicate representative VWF strings decorated with platelets. **C and D.** Immunoblot analysis and corresponding quantitative analysis, respectively, of platelet-associated TGF-β1 expression under no shear (control), shear, and shear plus supplementation of plasma-derived VWF. β-actin protein is used for the loading controls. **E.** Representative immunofluorescence images showing platelet CD61 (green) together with TGF-β1 (red) and PDGF-BB (magenta) following shear-induced activation. **F and G.** Flow cytometric plots and quantification P-selectin positive platelets, respectively, in *Vwf^+/+^*, *Vwf^+/-^* and *Vwf*⁻*^/^*⁻ rats under Sham vs. CTEPH conditions. **H**. Plasma TGF-β1 concentrations by ELISA in *Vwf^+/+^*, *Vwf^+/-^*and *Vwf*⁻*^/^*⁻ rats under Sham vs. CTEPH conditions. **I.** Representative immunohistochemical staining of lung sections showing TGF-β1 (top row), PDGF-B (2^nd^ row), endothelin-1 (3^rd^ row), and Ki-67 (bottom row) in pulmonary arterioles from *Vwf^+/+^*, *Vwf^+/-^* and *Vwf*⁻*^/^*⁻ rats under Sham vs. CTEPH conditions. **J** and **K.** Quantitative analysis of TGF-β1–positive area and PDGF-B–positive area, respectively, within pulmonary arterioles in *Vwf^+/+^*, *Vwf^+/-^* and *Vwf*⁻*^/^*⁻ rats following Sham vs. CTEPH procedure. Data presented are individuals (dots), means (bars) ± standard errors of the means (SEM) (horizontal lines). The differences were analyzed by Kruskal–Wallis test followed by Dunn’s multiple-comparisons test (**D**) or two-way ANOVA followed by the Tukey’s multiple comparisons test (**G, H, J and K**). P values less than 0.05 and 0.01 are significant and highly significant, respectively.

We further examined the downstream release of pro-remodeling mediators. While shear stress alone induced only a modest change in platelet-associated TGF-β1, the addition of UL-VWF under shear robustly enhanced platelet activation (**Figure S6A-C)** and subsequent TGF-β1 release (**Figure 6C-D, Figure S6D**). Imunofluorescence staining further confirmed colocalization of activated platelets with TGF-β1 and PDGF-BB (**Figure 6E**). Consistent with these in vitro findings, flow cytometric analyses demonstrated significantly reduced platelet activation in *Vwf*⁻*^/^*⁻ rats compared that in*Vwf^+/+^* rats after CTEPH induction (**Figure 6F-G**). In vivo, plasma TGF-β1 and PDGF-BB levels increased markedly in *Vwf^+/+^* rats following CTEPH induction, but significantly reduced in Vwf⁻/⁻ rats, with *Vwf^+/^*⁻ showing an intermediate phenotype (**Figure 6H**, **Figure S7A**).

Finally, reduced UL-VWF–dependent platelet activation was associated with diminished vascular accumulation of platelets and platelet-derived mediators. Compared with *Vwf^+/+^* rats, *Vwf*⁻*^/^*⁻ rats exhibited significantly lower CD61-positive area and reduced TGF-β1 and PDGF-BB deposition within pulmonary arteries, with *Vwf*⁺*^/^*⁻ rats again showing an intermediate phenotype (**Figure 6I–K, Figure S7B**). Notably, these changes were accompanied by attenuated endothelin-1 expression, a downstream effector of pro-remodeling signaling, and fewer Ki-67–positive proliferating cells within pulmonary arteries (**Figure S7C-D**). Collectively, these findings suggest that excessive UL-VWF–mediated platelet adhesion under shear promotes both thrombus formation and the release of platelet-derived pro-remodeling mediators, TGF-β1 and PDGF-BB, representing a critical mechanistic link between thrombosis and vascular remodeling in CTEPH.

### Recombinant ADAMTS13 ameliorates thrombosis and pulmonary vascular remodeling in CTEPH rats

Building on in vitro microfluidic thrombosis assays demonstrating that recombinant truncated ADAMTS13(rA13-T5, or rA13 in short) markedly suppresses VWF-mediated thrombus formation, we next evaluated whether rA13 exert similar antithrombotic and disease-modifying effects in vivo. WT rats subjected to the CTEPH procedure received either vehicle or rA13 (**Figure 7A, Figure S8A-B**). Compared to vehicle-treated controls, rA13-treated rats demonstrated enhanced survival rate (**Figure S8C**) and significantly improved pulmonary hemodynamics with lower RVSP, as reflected by increased PAT/PET ratios (**Figure 7B-E**). Structural remodeling and ventricular strain were also markedly attenuated, evidenced by reduced Fulton indices, RVWTD, and plasma NT-proBNP levels (**Figure 7F–G, Figure S8D**). Functionally, microfluidic assays showed that while CTEPH blood exhibited enhanced shear-induced thrombus formation, rA13 treatment significantly reduced both clot formation and stability (**Figure 7H-I**). Histological analyses further demonstrated prominent pulmonary arterial remodeling and in situ thrombosis in untreated CTEPH rats, characterized by increased vascular wall thickness, pulmonary microthrombi density, and accumulation of VWF- and CD61-positive material within pulmonary arteries. These pathological features were significantly attenuated following rA13 treatment (**Figure 7J and N–Q**), demonstrated by representative lung immunohistochemistry for fibrin, platelets, and VWF (**Figure 7R**). These improvements were accompanied by a normalized VWF multimer pattern and lower plasma D-dimer levels (**Figure 7K–M, Figure S8E**). Importantly, rA13 administration did not prolong tail bleeding time (**Figure S8F**). Together, these results indicate the preventive and therapeutic efficacy of recombinant ADAMTS13 in mitigating thrombus burden and pulmonary vascular remodeling in the CTEPH rats.

**Figure 7.**
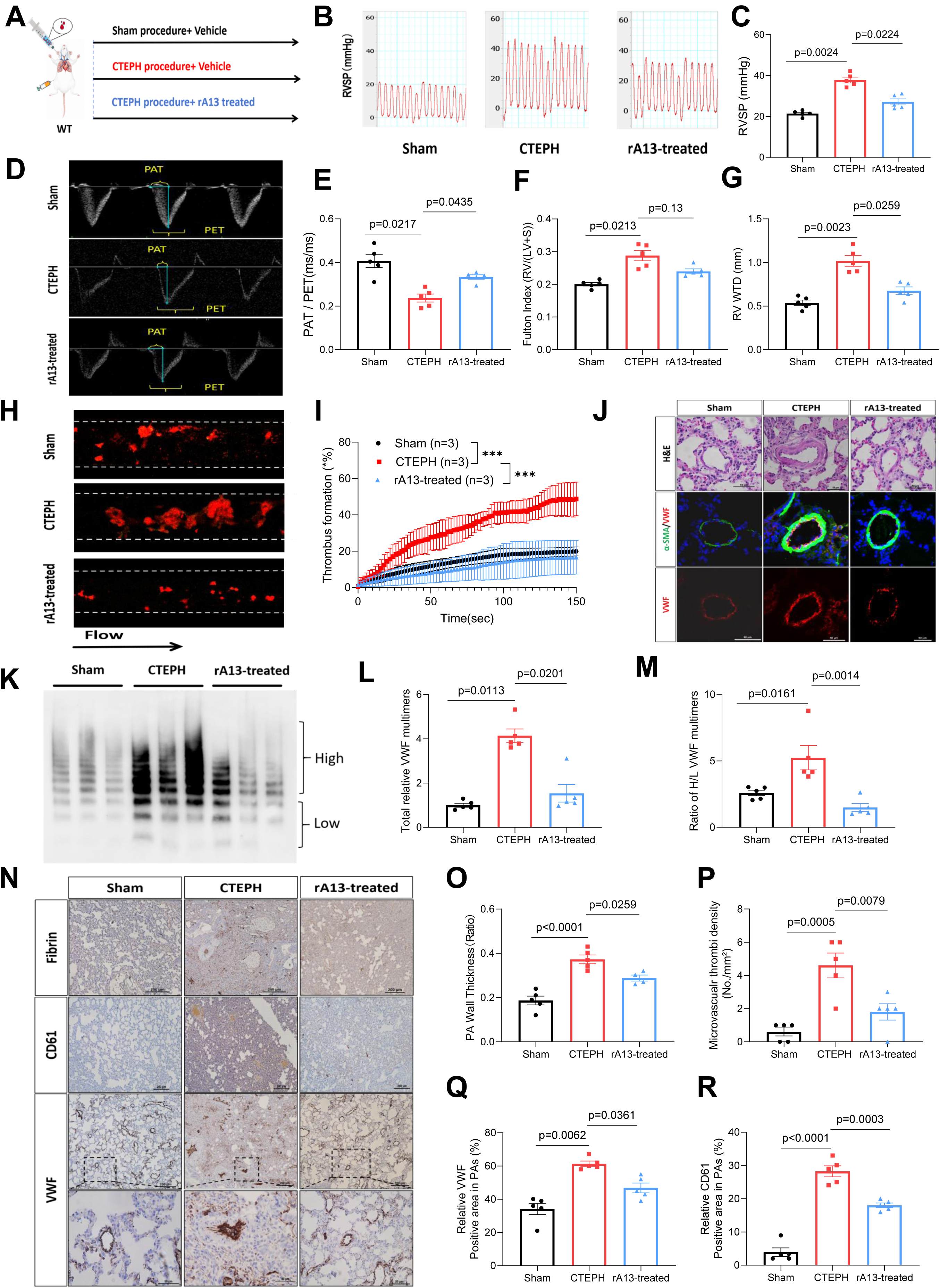
Recombinant ADAMTS13 reduces pulmonary thrombosis and vascular remodeling in CTEPH rats. **A.** Experimental design showing wild-type rats subjected to Sham or CTEPH procedures with vehicle or rA13 treatment. **B** and **C.** Representative waveforms and quantitative analysis of RVSP, respectively, in Sham, vs. CTEPH rats treated without or with rA13. **D** and **E.** Representative Doppler echocardiographic images and quantitative analysis of PAT/PET ratio, respectively in Sham and CTEPH rats treated without or with rA13. **F** and **G.** Fulton index and RVWTD, respectively, in Sham vs. CTEPH treated without or with rA13. **H** and **I.** Representative fluorescent images and quantitative analysis of kinetic thrombus formation over time, respectively, in the microfluidic shear-based assay of whole blood from Sham vs. CTEPH rats treated without or with rA13. **J.** Representative pulmonary arterial H&E staining (top row) and immunofluorescence staining for α-SMA (middle row) and VWF (bottom row) in Sham vs. CTEPH rats treated without or with rA13. **K, L,** and **M.** Plasma VWF multimer distribution, total VWF intensity, and ratios of high to low molecular weight VWF multimers, respectively, in Sham vs. CTEPH rats treated without or with rA13. **N.** Representative immunohistochemical images for fibrin (top row), CD61 (middle row), and VWF (bottom row) in the pulmonary arterioles from Sham vs. CTEPH rats treated without or with rA13. **O, P, Q,** and **R.** Pulmonary arterial wall thickness, pulmonary microthrombi density, area of pulmonary arteriole with positive VWF staining, and area of pulmonary arteriole with positive CD61 staining, respectively in Sham vs. CTEPH rats treated without or with rA13. Data shown are individual values (dots), means (bars) ± standard errors of the means (SEM) (horizontal lines). The differences were analyzed by one-way ANOVA followed by the Tukey’s multiple comparisons test. For time-course thrombosis analysis, two-way repeated-measures ANOVA was applied. P values less than 0.05 and 0.01 are significant and highly significant, respectively.

### Recombinant ADAMTS13 prevents platelets activation and platelet-associated pro-remodeling signaling in CTEPH rats

To assess if recombinant ADAMTS13 treatment attenuated VWF-dependent platelet activation and platelet-associated pro-remodeling signaling in vivo, platelet- and tissue-level analyses were performed. Flow cytometric analysis demonstrated increased platelet activation in CTEPH rats as assessed by P-selectin expression, whereas rA13 treatment reduced platelet activation (**Figure 8A**). Consistent with these observations, immunoblot analysis of washed platelet lysates revealed significantly increased platelet-derived TGF-β1 in CTEPH rats, including both latent (LAP-bound) and active dimeric forms, which were normalized following rA13 administration (**Figure 8B**-**C**). The rA13 treatment also significantly reduced platelet PDGF-BB levels compared with untreated CTEPH platelets (**Figure 8B and D**). In parallel, plasma ELISA measurements revealed concordant reductions in circulating TGF-β1 and PDGF-BB in rA13-treated group (**Figure 8E-F**), supporting an attenuation of platelet-derived pro-remodeling signaling with ADAMTS13 treatment.

**Figure 8.**
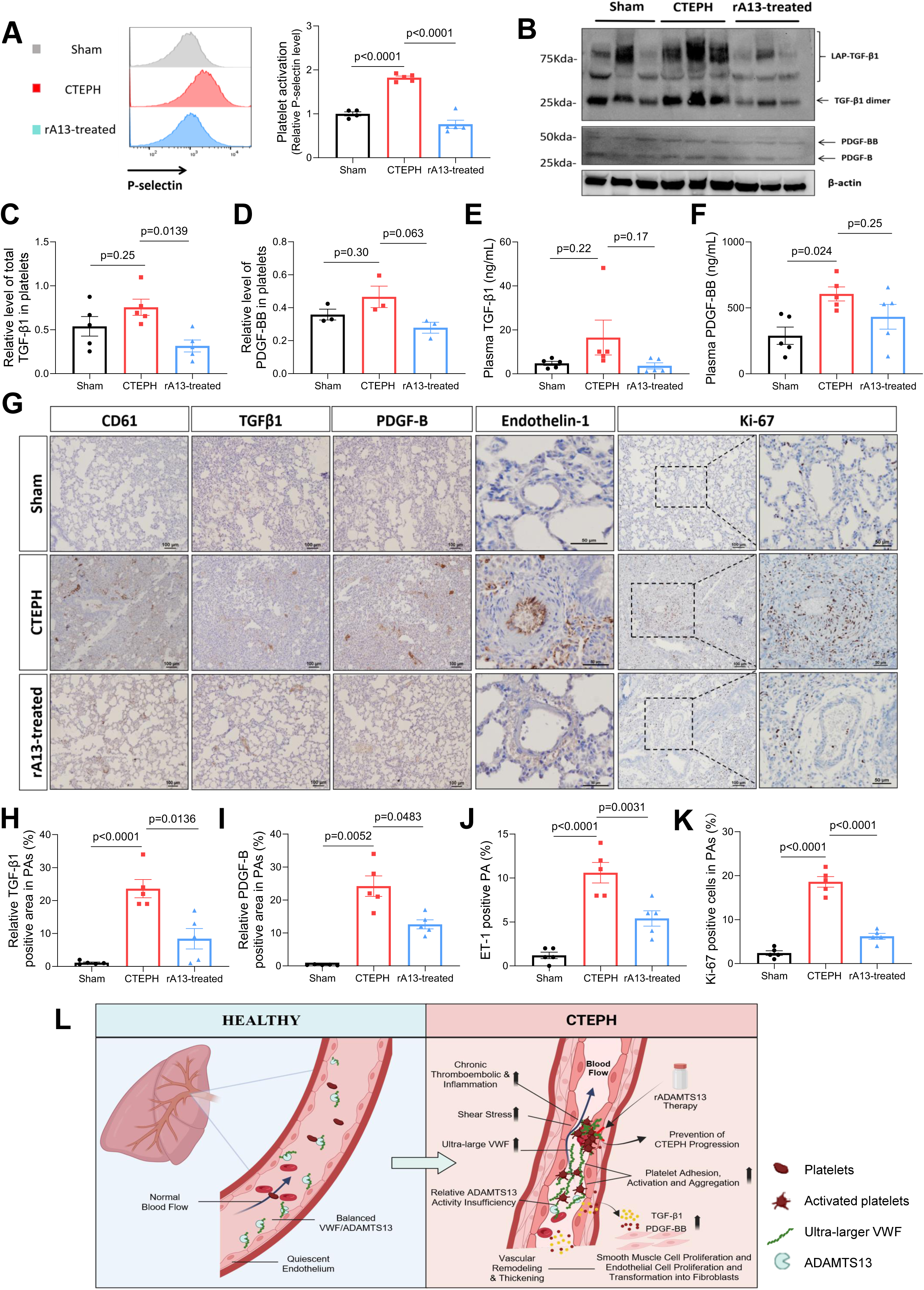
Recombinant ADAMTS13 suppresses platelet activation and the release of pro-remodeling factors in CTEPH rats. **A.** Flow cytometric profiles and relative P-selectin expression in Sham and CTEPH rats treated without or with rA13. **B.** Western blot analysis of platelet lysates showing TGF-β1 (latency-associated peptide, LAP and dimer forms) and PDGF-BB, with β-actin as a loading control. **C** and **D.** Relative levels of TGF-β1 and PDGF-BB in platelets of Sham vs. CTEPH rats treated without or with rA13. **E** and **F**. Plasma levels of TGF-β1 and PDGF-BB, respectively, in Sham vs. CTEPH rats treated without or with rA13, measured by ELISA. **G.** Representative immunohistochemical staining for CD61 (column 1), TGF-β1 (column 2), PDGF-B (column 3), endothelin-1 (column 4), and Ki-67 (column 5) in lung tissues from Sham vs. CTEPH rats treated without or with rA13. **H, I, J** and **K**. Quantification of TGF-β1, PDGF-B, Endothelin-1 and K--67 positive area, respectively, in pulmonary arteries of Sham vs. CTEPH rats treated without or with rA13. **L**. Schematic illustration depicting the proposed mechanism in pathogenesis of CTEPH, characterized by the relative deficiency of plasma ADAMTS13 activity and excessive accumulation of ultra-large VWF in the circulation and on pulmonary arterial endothelium, which promotes VWF–platelet adhesion, platelet activation and subsequent release of platelet-derived pro-remodeling mediators, such as TGF-β1 and PDGF-BB, which collectively contribute to pulmonary vascular remodeling. In contrast, recombinant ADAMTS13 treatment restores the ADAMTS13/VWF balance and modulate platelet-driven pulmonary vascular remodeling. The schematic was generated using BioRender.com. Data presented are the mean ± SEM. The differences were analyzed by one-way ANOVA followed by the Tukey’s test. P values are indicated in individual panels.

To determine the effects of platelet-driven pro-remodeling factors on the pulmonary vasculature, lung immunohistochemical assessment was performed. As shown **in Figure 8 G-I**, CD61-positive platelets accumulation, TGF-β1, and PDGF-B levels markedly increased within pulmonary arteries of CTEPH rats, which were largely normalized following rA13 treatment. Elevated Endothelin-1 and the proliferation marker Ki-67 positive cells, which indicate the vascular remodeling in CTEPH lung, were also decreased following rA13 treatment (**Figures 8G, J**, and **K**). Taken together, these findings support a mechanism in which insufficient ADAMTS13 activity, coupled with accumulation of ultra-large VWF in the circulation and on pulmonary arterial endothelium, promotes persistent thrombus formation and platelet activation. Activated platelets subsequently release pro-remodeling mediators, including TGF-β1 and PDGF-BB, thereby driving pulmonary vascular remodeling. Genetic ablation of VWF or therapeutic restoration of ADAMTS13 activity suppresses thrombosis and platelet activation and attenuates platelet-driven pulmonary vascular remodeling as schematically outlined in **Figure 8L**.

## Discussion

Here, using human CTEPH samples and new rat models, we identify the dysregulation of VWF-ADAMTS13 axis as a pathogenic drive of CTEPH, linking dysregulated thrombosis and platelet adhesion/activation to downstream endothelial dysfunction and pulmonary arterial remodeling. Our results further demonstrate that UL-VWF–driven platelet activation and subsequent release of platelet-driven pro-remodeling factor, including TGF-β1 and PDGF-BB, represent an important mechanism by which thrombotic events are transduced into enhanced pulmonary vascular remodeling process. Most importantly, the lack of VWF or restoration of ADAMTS13 function efficiently disrupts this pathogenic cascade in the rat models of CTEPH, which provides basis for targeted therapy.

Modeling CTEPH in murine animals is challenging, largely because their highly efficient fibrinolytic system rapidly clears pulmonary emboli, preventing the persistent, organized vascular obstructions that recapitulate the human disease(29, 30). In our study, the autologous clot injection combined with SU5416 represents a superior rat model of CTEPH because it uniquely integrates the two central drivers of human disease: persistent thromboembolic obstruction and impaired vascular repair. Repeated administration of autologous clots mimics the clinical origin of CTEPH from unresolved pulmonary emboli, enabling emboli to lodge, organize, and resist fibrinolysis, while avoiding immunogenic artifacts associated with injection of heterologous clots or synthetic materials. The addition of SU5416, a VEGFR2 inhibitor, suppresses endothelial regeneration and promotes a pro-remodeling environment, thereby facilitating thrombus persistence and progressive vascular remodeling(30, 31). This combined approach yields stable pulmonary hypertension, right ventricular hypertrophy, microvascular thrombosis, and arterial remodeling that closely recapitulate human CTEPH.

Platelet-VWF interaction is important for thrombosis and inflammation. Platelet glycoprotein 1b (GP1b)(32, 33) is the receptor for VWF that is released from vascular endothelium upon stimulation by inflammatory cytokines or thrombin(34–36). This process is tightly regulated by the circulating or locally produced ADAMTS13 that cleaves the Tyr-Met bond in the central A2 domain of VWF(37, 38). Thus, ADAMTS13 functions as a key safeguard that preserves vascular homeostasis and restrains thrombus formation and inflammation(39, 40). In the context of CTEPH, the marked accumulation of UL-VWF in circulation and in situ shifts this balance towards relative deficiency of ADAMTS13 activity, which is unable to counteract the increased UL-VWF-platelet interaction and aggregation. In our rat CTEPH model, we found that ADAMTS13 deficiency further exacerbates pulmonary microvascular thrombosis, resulting in early mortality. However, once the rats survived initial PE, the remaining *Adamts13^-/-^* rats did not exhibit more exacerbated hemodynamic burden than WT rats following CTEPH induction. This may be confounded by the significant elevation of VWF expression and reduction of ADAMTS13 in WT rats following CTEPH induction, rendering severe deficiency of VWF proteolysis in both WT and *Adamts13^-/-^*rats. Clearly, addition of recombinant ADAMTS13 significantly reduces the early mortality, pulmonary arteriole thrombosis, and vascular remodeling following CTEPH induction, supporting the protective role of ADAMTS13 in PE and CTEPH.

The molecular mechanism underlying ULVWF-platelet enhanced vascular remodeling is not fully understood. Our results demonstrate the upregulation and release of PDGF-BB and TGF-β1 that have been shown to promote pulmonary arterial remodeling in pulmonary hypertension(41–44) may contribute to progressive vascular remodeling in the CTEPH rats through endothelial activation and endothelin-1 production(45). Similar increase levels of these remodeling factors were observed in patients with CTEPH(45). Platelets serve as a major reservoir of TGF-β1 and PDGF-BB(46–48). Under shear condition, endothelial UL-VWF–dependent platelet adhesion robustly enhances platelet activation, triggering the coordinated release of platelet-stored pro-remodeling mediators within the pulmonary arteries. This platelet-derived signaling converges on the pulmonary endothelium and vascular wall, establishing a mechanistic link between thrombosis, impaired thrombus resolution, and vascular remodeling.

Several therapeutic strategies that target UL-VWF–platelet interaction are available today. This includes the agents that directly block UL-VWF and platelet binding such as anti-VWF A1 nanobody (caplacizumab)(49–51) and anti-GP1b antagonist (anfibatide)(52, 53) or humanized monoclonal antibody against GP1bα(54). In addition, recombinant ADAMTS13 is now Food and Drug Administration-approved drug for treatment of TTP(55). Among these therapeutic approaches, we focused on the efficacy of recombinant ADAMTS13 because it restores a physiological regulatory mechanism governing VWF multimer size, thereby dismantling UL-VWF–platelet interactions without causing bleeding. The therapeutic efficacy of recombinant ADAMTS13 in our rat CTEPH model highlights that UL-VWF–driven platelet activation may represent a pathogenic pathway not adequately targeted by conventional anticoagulant or thrombolytic therapies.

There are limitations of this study. Human plasma samples were collected using EDTA, precluding the assessment of plasma ADAMTS13 activity. In addition, while we focused on platelet-derived pro-remodeling mediators, activated platelets may release a broad spectrum of bioactive factors that may also contribute to pulmonary vascular remodeling and need to be investigated in future study.

Nevertheless, we conclude that our study demonstrates the direct pathogenic roles of excessive VWF and ADAMTS13 deficiency in the development of CTEPH. We also demonstrate that UL-VWF–mediated platelet adhesion and activation is a key upstream driver for releasing pro-remodeling mediators, which is mechanistically linked to pulmonary microvascular thrombosis, impaired thrombus resolution, and subsequent development of CTEPH. Targeting the dysregulation of VWF-ADAMTS13 axis may represent a promising therapeutic strategy to prevent persistent thrombosis and to attenuate pulmonary vascular remodeling in CTEPH and other pulmonary thromboembolic diseases.

## Non-conventional Abbreviation

ADAMTS13: A disintegrin and metalloproteinase with thrombospondin type 1 repeats member 13
CTEPH: chronic thromboembolic pulmonary hypertension
DMSO: dimethyl sulfoxide
EDTA: ethylenediaminetetraacetic acid
ELISA: enzyme-linked immunosorbent assay
FRETS: fluorescence resonance energy transfers
GP1b: glycoprotein Ib
IPAH: idiopathic pulmonary arterial hypertension
PAT/PET: Pulmonary acceleration time/pulmonary ejection time
PDGF-BB: platelet-derived growth factor BB homodimer
PE: pulmonary embolism
PH: pulmonary hypertension
PTE: pulmonary thromboendarterectomy
PVDF: polyvinylidene fluoride
RVSP: right ventricular systolic pressure
SDS-PAGE: sodium dodecyl sulfate polyacrylamide gel electrophoresis
SU5416: a potent and selective inhibitor of the VEGFR2 receptor
TGF-β1: transforming growth factor beta-1
TMA: trimethylamine
TTP: thrombotic thrombocytopenic purpura
UL-VWF: ultra large von Willebrand factor
VWF: von Willebrand factor
VWF73 or VWF71: a 73 or 71 amino acid peptide derived from central A2 domain of VWF

## Authors’ contributions

Z.W, L.Z. and X.L.Z. conceived the research. Z.W, L.Z., H.D., Q.Z., Z.C. and A.L. performed experiments and collected data. Z.W., L.Z. and X.L.Z. analyzed and interpreted data. Z.W. and L.Z. wrote the manuscript. L.Z and X.L.Z revised the manuscript. L.S., X.Z, E.M.D. and M.J.S. provide key experimental materials. All authors approved the final version of the manuscript for submission.

## Acknowledgments

We thank Dr. Jian Huang of the Cardiovascular Research Core, University of Kansas Medical Center (KUMC) for assistance with transthoracic echocardiography in rats, Dr. Navneet K. Dhillon, Department of Internal Medicine, KUMC for assistance with right ventricular systolic pressure measurement during the pilot phase of model establishment, and Lily He of the Pathology Core, KUMC for expert assistance with rat histopathological analyses. We also thank Dr. Joshua Muia of the Versiti Blood Research Institute for providing the cattle-derived FRETS-VWF73 substrate and Drs. Robert R. Montgomery and Christian Kastrup for providing *Vwf*-deficient rat.

## Supplemental Materials

**Figure S1.**
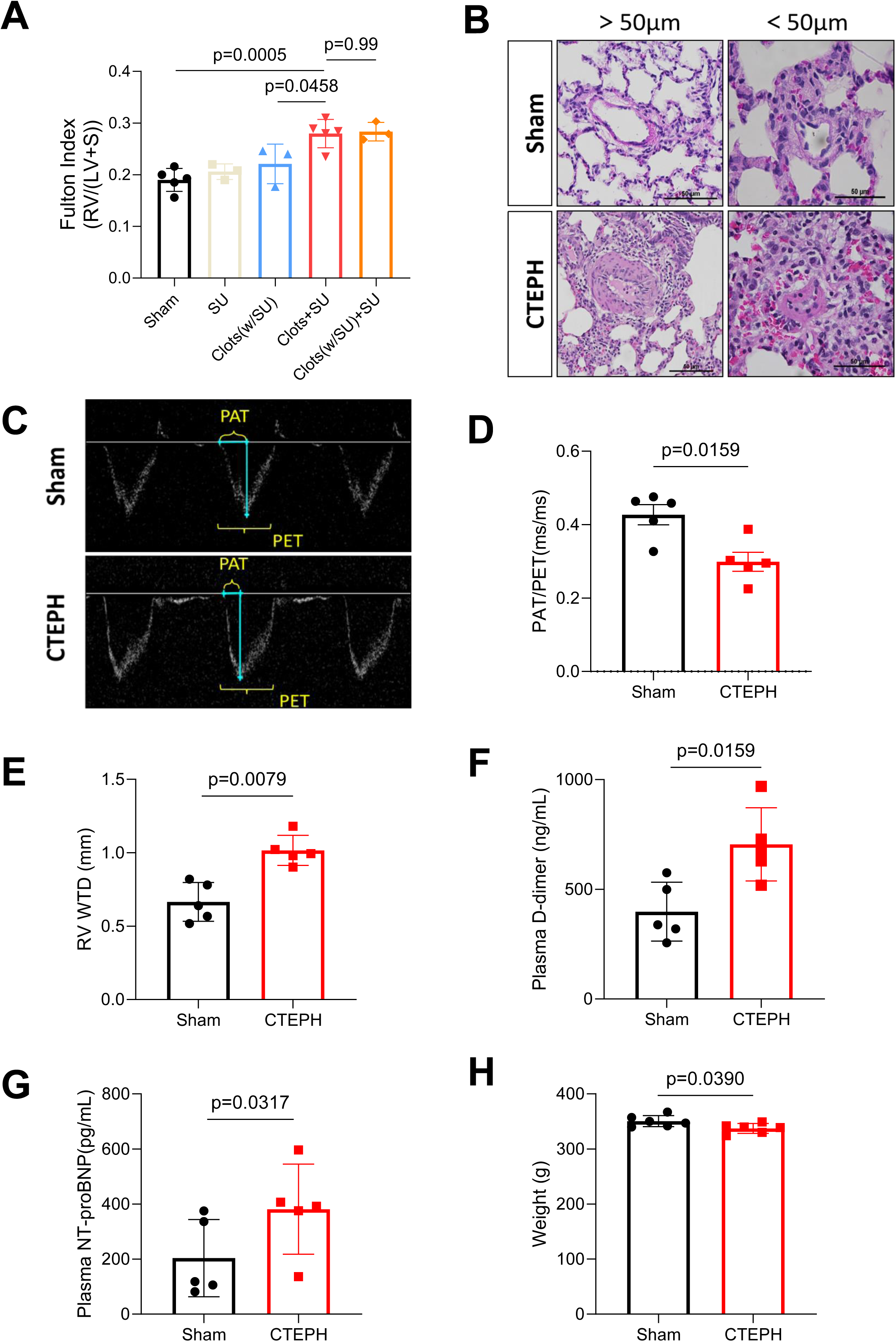
Supplementary phenotypic validation of the optimized rat CTEPH model. **A.** Fulton index [RV/(LV+S)] measured across the five experimental groups. **B.** H&E staining showing the remodeling of pulmonary arteries (PAs) with diameters larger than 50 μm and smaller than 50 μm. **C** and **D.** Representative Doppler echocardiographic images of pulmonary artery flow showing pulmonary artery acceleration time (PAT) and pulmonary ejection time (PET) and the quantification of the PAT/PET ratio, respectively. **E.** Echocardiographic measurement of right ventricular wall thickness during diastole (RVWTD). **F.** Plasma D-dimer levels measured at the terminal time point. **G.** Plasma NT-proBNP levels in Sham vs. CTEPH rats. **H.** Body weight comparison between Sham vs. CTEPH rats. Data presented are the mean ± SEM and the differences were analyzed by one-way ANOVA followed by the Tukey’s multiple comparisons test for comparison among three groups or two-tailed unpaired t-test for two groups.

**Figure S2.**
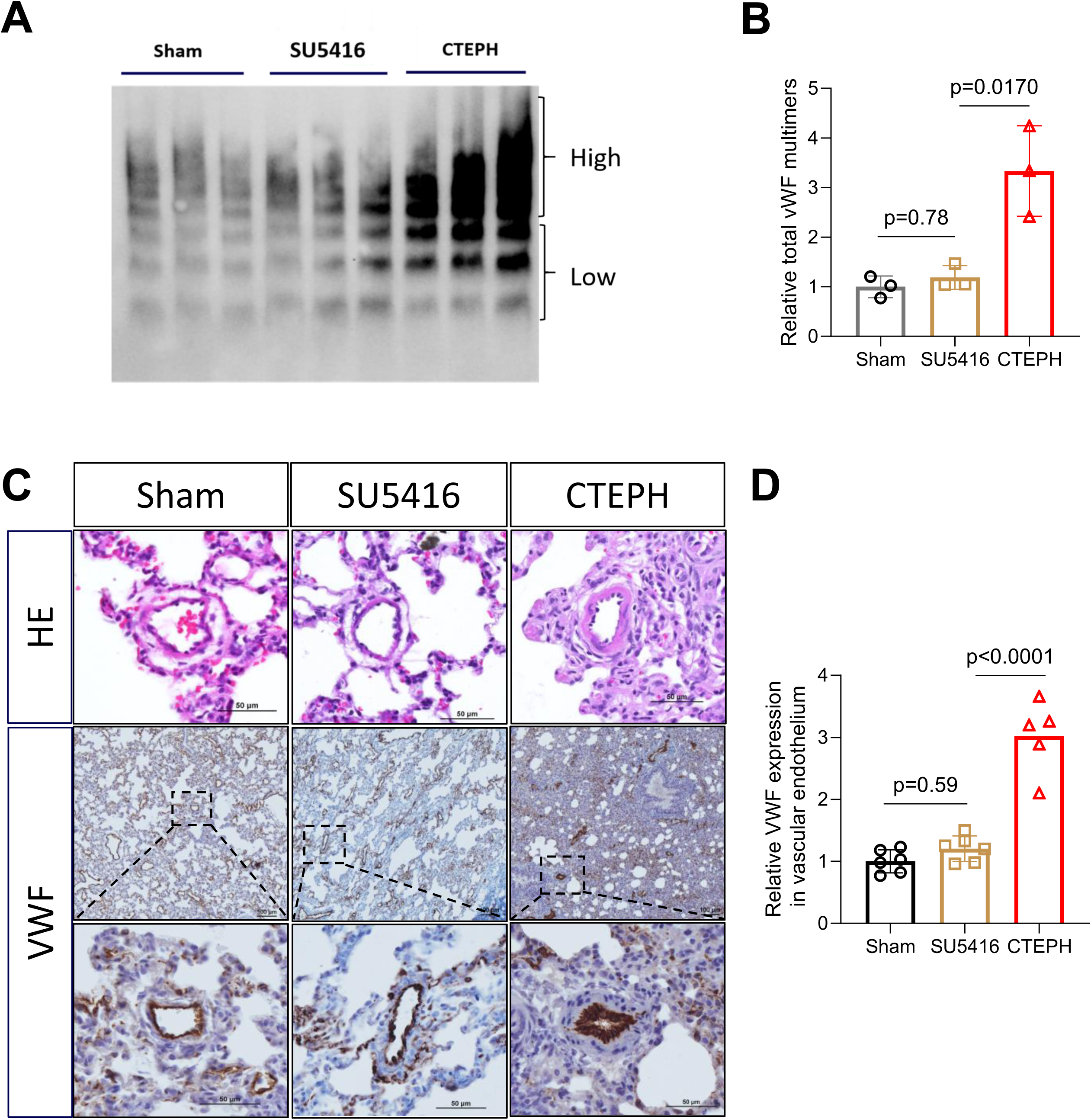
Subcutaneous SU5416 administration does not alter plasma VWF multimers and pulmonary endothelial VWF expression in rats. **A.** Representative agarose gel electrophoresis showing plasma VWF multimer distribution from rats received Sham, SU5416 alone, and CTEPH protocol. **B.** Quantification of total relative plasma VWF multimers. **C.** Representative H&E staining and immunohistochemical staining for VWF in the lung tissue sections from Sham, SU5416, and CTEPH rats. Dashed boxes indicate regions shown at higher magnification. **D.** Quantification of relative VWF expression in the pulmonary vascular endothelium. Data presented are the mean ± SEM and the differences were analyzed by one-way ANOVA followed by the Tukey’s multiple comparisons test.

**Figure S3.**
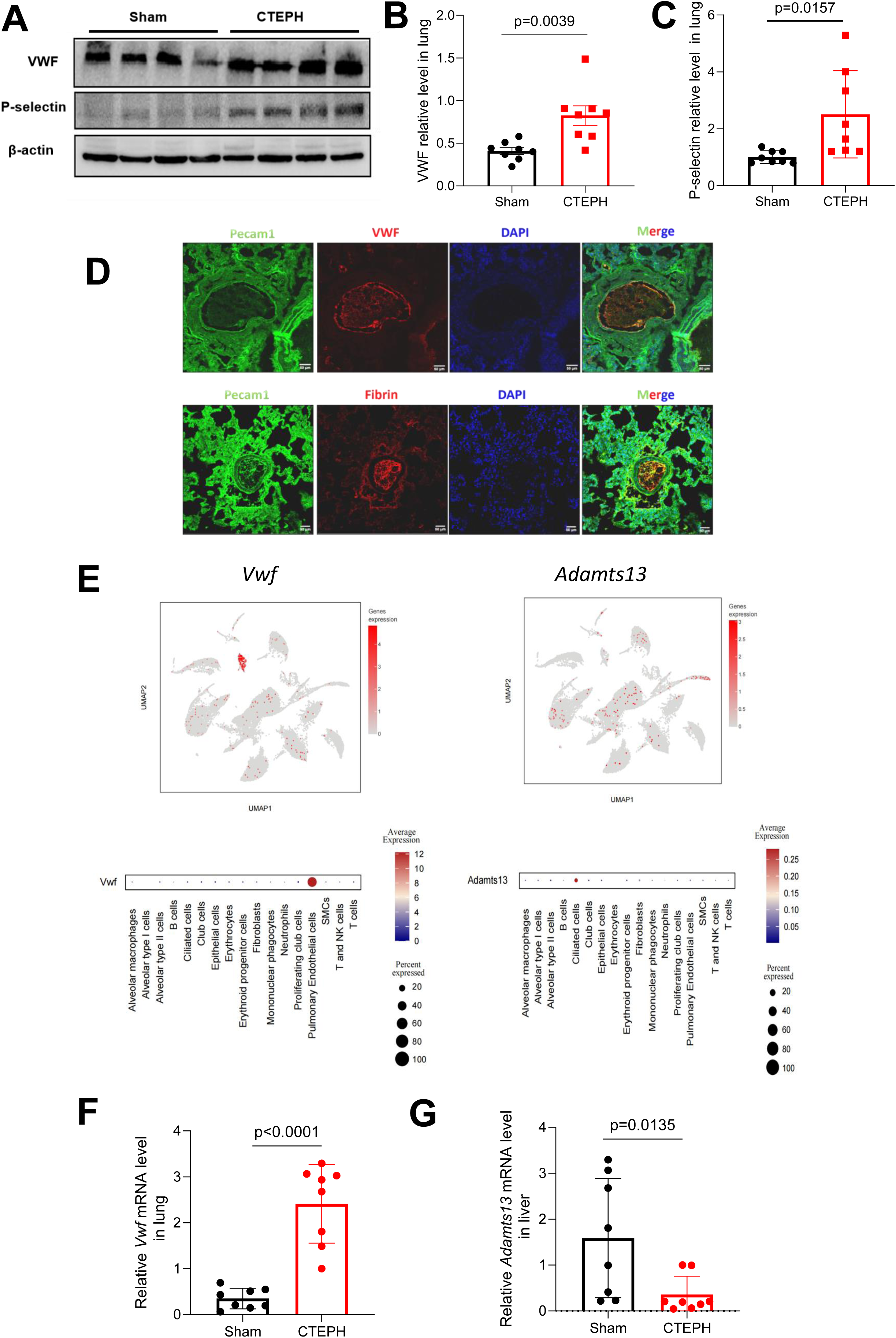
Increased pulmonary VWF expression but reduced hepatic expression of *Adamts13* mRNA in CTEPH rats. **A**. Western blot analysis of total VWF and P-selectin protein levels in whole lung lysates from Sham vs. CTEPH rats. **B** and **C**. Densitometric quantification of lung VWF and P-selectin protein levels, respectively, following normalization to β-actin. **D**. Representative immunofluorescence staining of pulmonary microthrombi showing fibrin and VWF expression. **E**. Featured plots from single-cell RNA sequencing showing the cellular distribution of *Vwf* and *Adamts13* mRNA across lung cell populations. **F**. Quantitative real-time PCR validation of VWF mRNA expression in lung tissue from Sham vs. CTEPH rats. **G**. Quantitative real-time PCR analysis of *Adamts13* mRNA expression in liver tissue from Sham vs. CTEPH rats. Data are presented as mean ± SEM. Comparisons between two groups were performed using unpaired two-tailed Student’s t test.

**Figure S4.**
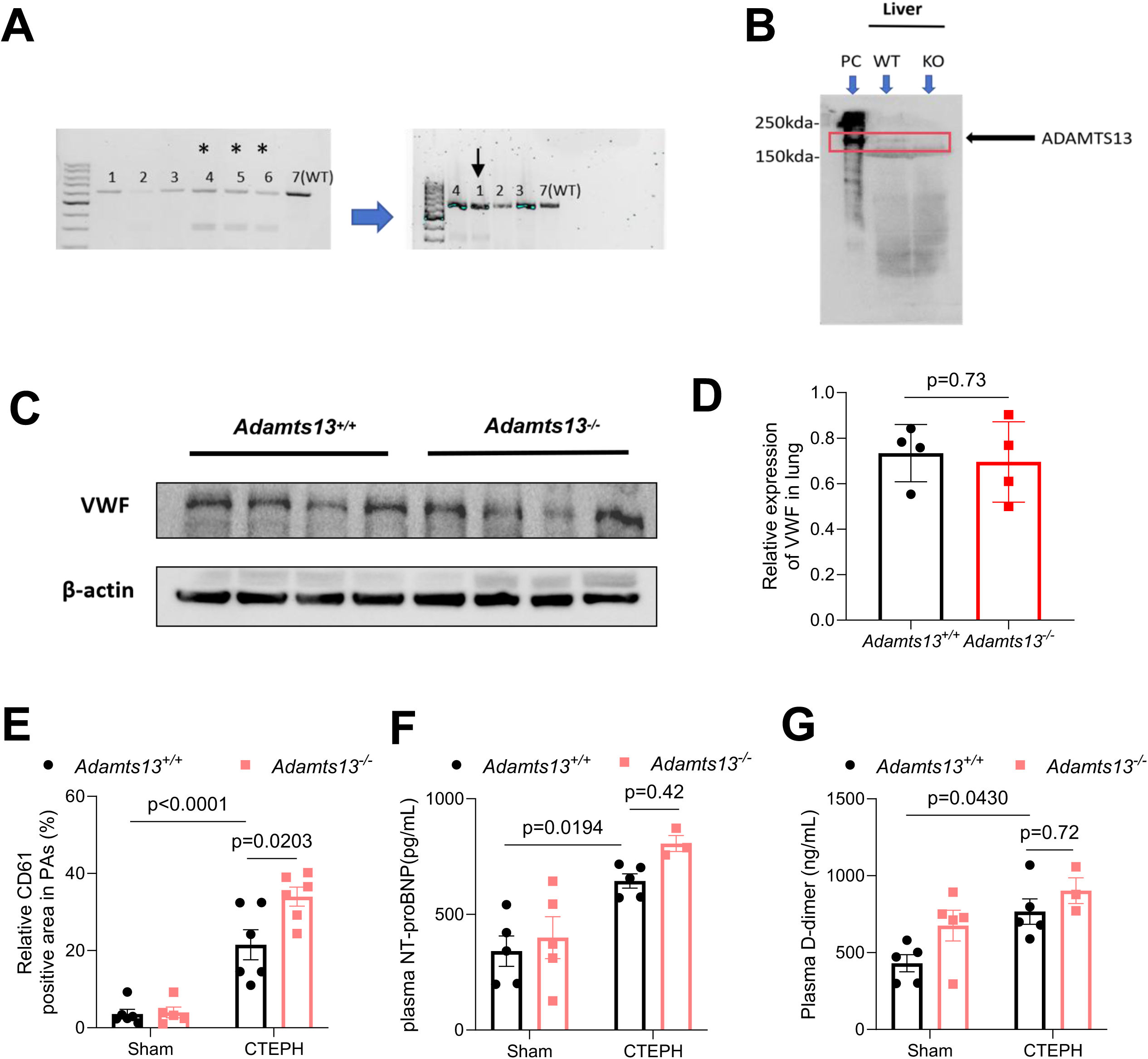
Validation of *Adamts13* knockout and characterization of rats under Sham and CTEPH conditions. **A.** Representative agarose gel electrophoresis showing T7E1 endonuclease digestion of PCR products used for genotyping *Adamts13* rats. Lanes 4 and 5 correspond to *Adamts13* heterozygous rats, whereas lane 7 represents a wild-type (*Adamts13^+/+^*) rat. Following T7E1 digestion, PCR products were re-analyzed to distinguish genotypes. Lane 1 was identified as a homozygous Adamts13 knockout (*Adamts13^-/-^*) rat, while lanes 2 and 3 correspond to *Adamts13*⁺*/*⁺ rats. **B**. Immunoblot analysis of ADAMTS13 protein expression in liver tissue from *Adamts13^+/+^*and *Adamts13^-/-^* rats. PC, positive control; KO, *Adamts13^-/-^*. **C**. Representative immunoblot of VWF protein expression in lung tissue from *Adamts13^+/+^* and *Adamts13^-/-^* rats. **D**. Quantification of relative pulmonary arteries (PAs) VWF expression normalized to β-actin. **E**. Quantification of pulmonary CD61-positive area in PAs from *Adamts13^+/+^* and *Adamts13^-/-^* rats under Sham vs. CTEPH procedure. **F**. Plasma NT-proBNP levels in *Adamts13^+/+^*and *Adamts13^-/-^* rats under Sham vs. CTEPH conditions. **G**. Plasma D-dimer levels in *Adamts13^+/+^* and *Adamts13^-/-^*rats under Sham vs. CTEPH conditions. Data presented are the mean ± SEM and the differences were analyzed by two-tailed unpaired t-test **(D)** or two-way ANOVA followed by the Tukey’s multiple comparisons test (**E-G**).

**Figure S5.**
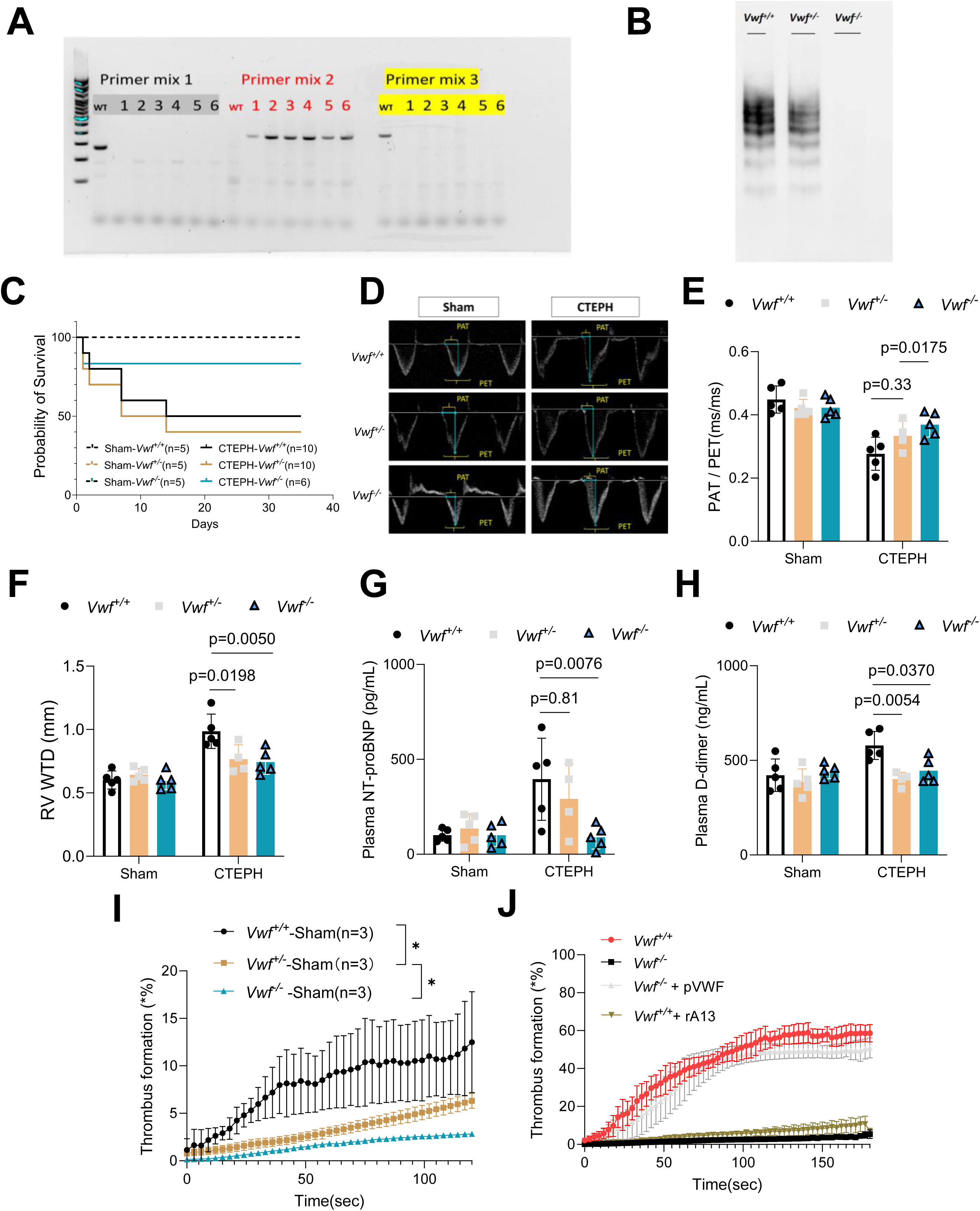
Genotyping, hemodynamic assessment, survival, thrombosis, and circulating biomarkers in *Vwf^-/-^*rats subjected to CTEPH procedure. **A.** Schematic illustration of PCR primer design and representative genotyping results used to identify *Vwf^+/+^*, *Vwf^+/-^* and *Vwf*⁻*^/^*⁻ rats. Primer sets distinguishing wild-type, deletion, and 5’-end alleles are shown together with representative agarose gel electrophoresis images. **B**. Representative plasma VWF multimer patterns from *Vwf^+/+^*, *Vwf^+/-^*and *Vwf*⁻*^/^*⁻ rats. **C**. Kaplan–Meier survival analysis of *Vwf^+/+^*, *Vwf^+/-^* and *Vwf*⁻*^/^*⁻ rats following Sham vs. CTEPH procedure. **D**. Representative Doppler echocardiographic images showing PAT/PET measurements in *Vwf^+/+^*, *Vwf^+/-^* and *Vwf*⁻*^/^*⁻ rats under Sham and CTEPH conditions. **E** and **F**. Quantitative analysis of PAT/PET ratio and RVWTD, respectively, in *Vwf^+/+^*, *Vwf^+/-^* and *Vwf*⁻*^/^*⁻ rats subjected to Sham vs. CTEPH procedure. **G**. Plasma NT-proBNP concentrations measured in *Vwf^+/+^*, *Vwf^+/-^* and *Vwf*⁻*^/^*⁻ rats under Sham vs. CTEPH conditions. **H**. Plasma D-dimer concentrations measured in *Vwf^+/+^*, *Vwf^+/-^* and *Vwf*⁻*^/^*⁻ rats under Sham vs. CTEPH conditions. **I**. Time-course quantification of thrombus formation in a microfluidic shear-based thrombosis assay using whole blood from *Vwf^+/+^*, *Vwf^+/-^* and *Vwf*⁻*^/^*⁻ rats under Sham conditions. **J**. Time-course analysis of thrombus formation in the microfluidic thrombosis assay using whole blood from *Vwf^+/+^* rats, *Vwf*⁻*^/^*⁻ rats, *Vwf*⁻*/*⁻ supplemented with purified plasma VWF (pVWF), and *Vwf^+/+^*blood treated with recombinant truncated ADAMTS13 (rA13). Data presented are the mean ± SEM. Differences were analyzed by two-way ANOVA followed by Tukey’s multiple comparisons test. For time-course thrombosis analyses, two-way repeated-measures ANOVA was applied.

**Figure S6.**
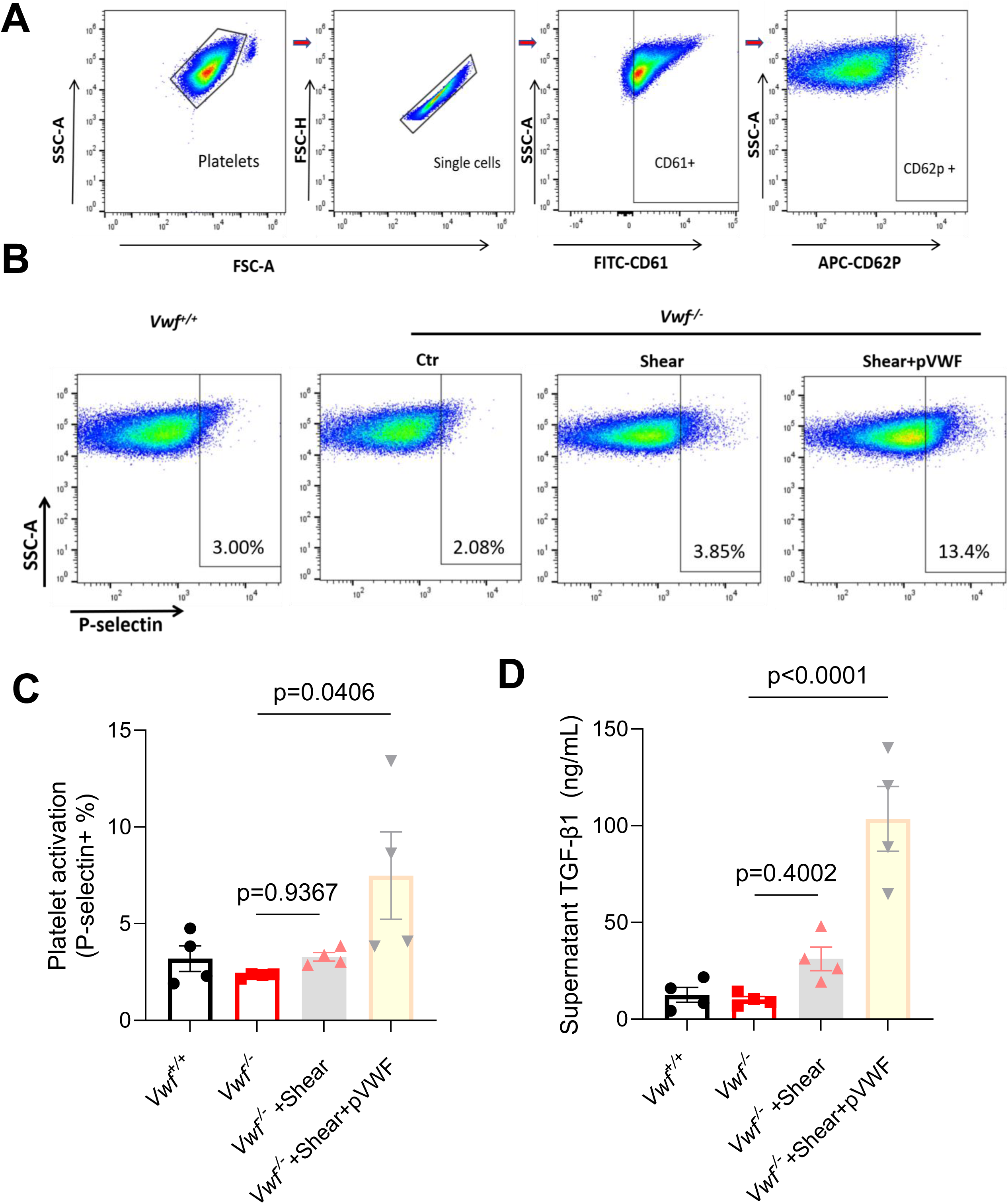
VWF-dependent platelet activation under shear. **A**. Gating strategy for flow cytometric analysis of platelet activation. Platelets were identified based on SSC-A/FSC-A properties, followed by singlet selection. CD61-positive platelet populations were gated and assessed for CD62P (P-selectin) expression as a marker of platelet activation. **B**. Representative flow cytometry plots showing P-selectin expression in platelets isolated from *Vwf^+/+^* and *Vwf*^⁻^*^/^*^⁻^ conditions under different experimental stimuli, including shear stress (Shear) and shear stress with added purified VWF (Shear+pVWF). Percentages of CD62P-positive platelets are indicated in each panel. **C**. Quantification of platelet activation (CD62P positivity) across experimental groups under different conditions. **D**. ELISA measurement of supernatant TGF-β1 levels (pg/mL) in the corresponding experimental conditions. Data are shown as individual values (dots), means (bars), and SEM (error bars). Statistical comparisons were performed using one-way ANOVA followed by appropriate post hoc tests. P values <0.05 were considered statistically significant.

**Figure S7.**
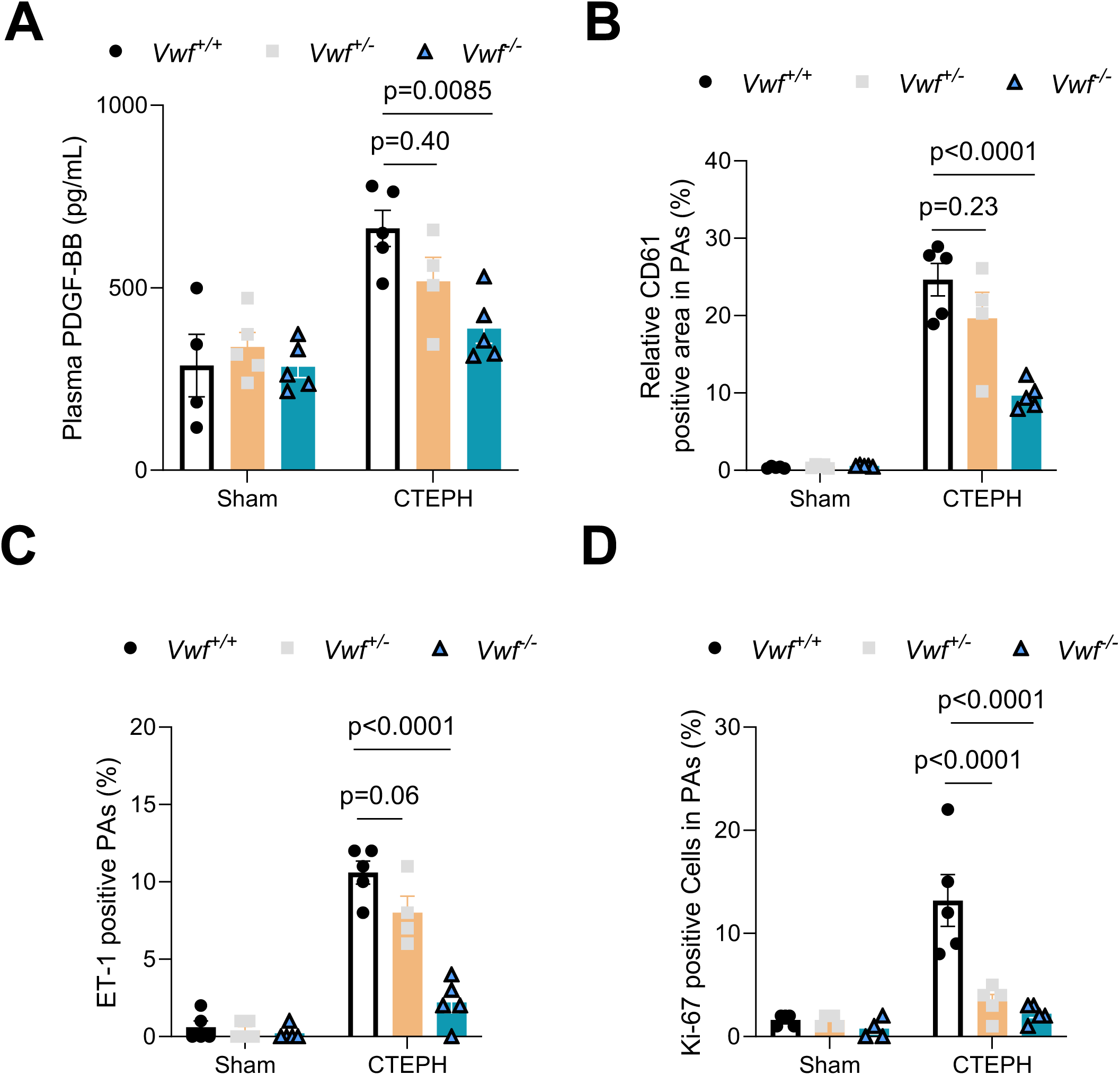
Genetic deletion of *Vwf* mitigates thromboinflammatory signaling, platelet vascular deposition, and arterial remodeling following chronic embolic challenge. **A.** Plasma PDGF-BB concentrations measured in *Vwf^+/+^*, *Vwf^+/-^*and *Vwf*⁻*^/^*⁻ rats under Sham vs. CTEPH conditions. **B, C**, and **D**. Quantification of platelet CD61-positive area, endothelin-1 (ET-1)–positive PAs, and Ki-67–positive cells within PAs, respectively, in lung sections from *Vwf^+/+^*, *Vwf^+/-^* and *Vwf*⁻*^/^*⁻ rats subjected to Sham vs. CTEPH procedure. Data presented are the mean ± SEM. Differences were analyzed by two-way ANOVA followed by Tukey’s multiple comparisons test. For time-course thrombosis analyses, two-way repeated-measures ANOVA was applied.

**Figure S8.**
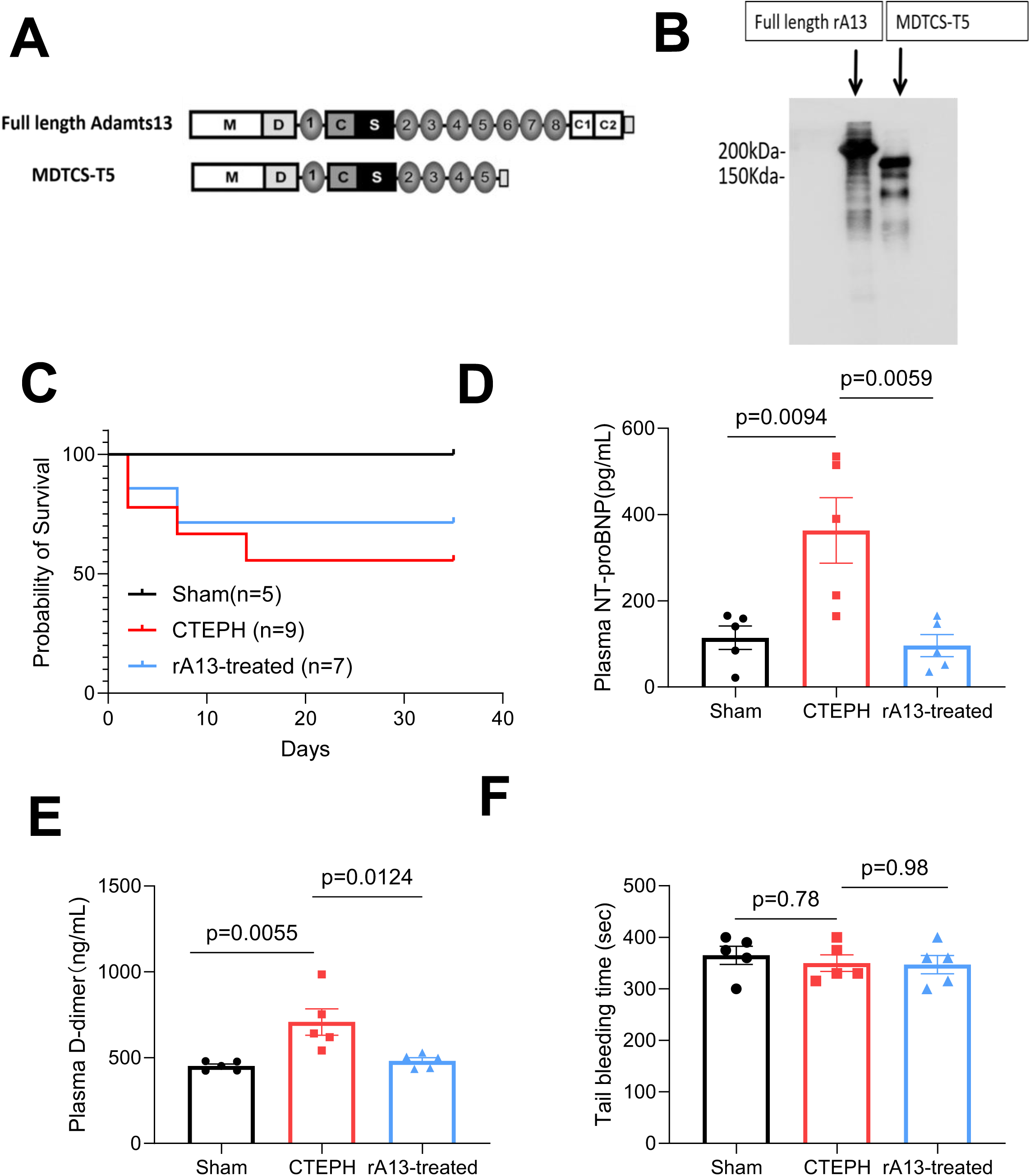
Characterization and systemic effects of recombinant ADAMTS13 in the rat CTEPH model. **A.** Schematic representation of the domain organization of full-length ADAMTS13 and the recombinant ADAMTS13 variant truncated after the 5^th^ TSP1 repeat (rA13-T5, or rA13 in short) used in this study. **B.** Representative immunoblot showing recombinant full-length ADAMTS13 and rA13-T5 proteins. **C.** Tail bleeding time measured in Sham, CTEPH, and rA13-treated CTEPH rats. **D.** Kaplan–Meier survival analysis of Sham, CTEPH, and rA13-treated CTEPH rats. **E.** Plasma NT-proBNP levels measured by ELISA. **F.** Plasma D-dimer levels measured by ELISA. Data presented are the mean ± SEM and the differences were analyzed by one-way ANOVA followed by the Tukey’s multiple comparisons test. P values are indicated in individual panels,

