## Supplementary materials for "Targeting pathogenic VWF/ADAMTS13 dysregulation attenuates CTEPH progression"

**Supplemental Materials**

**Methods**

##### *Rat genotyping*: This protocol describes genotyping of rat models using ear tissue collected before weaning (at age of 21 days). PCR products for genotyping were generated using Platinum Direct PCR Universal Master Mix kit (Thermo Fisher Scientific, Cat. #A44647100, MA, USA). A small piece of ear tissue was collected and placed in a microcentrifuge tube containing a lysis buffer and proteinase K. The tissues was incubated at room temperature for 10 minutes for lysis, then heated at 95°C for 1 minute to denature proteins and inactivate proteinase K. The supernatant was used directly as DNA template for PCR reaction using specific primers for the target genes and optimized thermal cycling conditions. Successful amplification of the PCR products was verified with 1.2% agarose gel electrophoresis. For the identification of *Adamts13* knockout rats, the PCR products were subjected to T7E1 digestion（New England Biolabs, Cat.#M0302S, MA, USA) per the manufacturer’s instructions^1^. The digested products were analyzed by agarose gel electrophoresis. If there were two bands, it indicated a heterozygous genotype; if it was a single band, it suggested either a homozygous genotype (knockout) or wild type because T7E1 cuts only heteroduplex, but not homoduplex. To confirm the genotype, the PCR products were also sent out for Sanger sequencing, which separated wild type (WT) from homozygous knockouts. Additional validation was performed by mixing the unknown PCR products with a known WT PCR products and perform T7E1 enzyme digestion. A single band following digestion indicates a WT in the unknown product, while two bands confirm the homozygous genotype in the unknown. All genotyping results of *Adamts13* knockout can be found in Supplementary Figure S3.

##### This protocol provides a quick and reliable workflow for identifying heterozygous, wild type, and homozygous knockout genotypes, particularly with those of point mutations or small deletions or insertions^2^. For identifying VWF heterozygous, knockout, and wild-type rats, we used three primer combinations (provided by the laboratory of Dr. Robert Montgomery at Versiti). The primer sets are listed in Supplementary Table S2. In wild-type rats, primers (Mix1 and Mix3) produce PCR bands; In heterozygous rats, primers (Mix1, Mix2, and Mix3) all yield bands; and in knockout rats, only primers (Mix2) generate a band. The PCR band patterns and the corresponding VWF multimer–based genotype validation are shown in Supplementary Figure S4A-B. All primer sequences for RT-QPCR are detailed in the Supplementary Table S1.

*Clot preparation*: Male Sprague-Dawley (SD) rats of different genotypes, aged 6 weeks and weighing 180–220 g, were used. Blood was collected from the orbital vein using a non-heparinized capillary tube (Globe Scientific, Cat. #51602, untreated, blue tip, USA). To generate SU5416-containing autologous thrombi, 0.5 μg of SU5416 dissolved in 50 μL DMSO was added to 950 μL of freshly collected whole blood, followed by rapid mixing to achieve a final concentration of 0.5 μg/mL SU5416. The blood mixture was immediately aspirated into non-heparinized capillary tubes and allowed to clot for subsequent thrombus preparation. Once coagulated, the blood clots were placed in 10% tranexamic acid (Cayman Chemical Company, Cat. #19193, USA) saline solution and stored at 4°C.

*Establishment of a rat model of CTEPH*: To develop a physiologically relevant rat model of CTEPH, we modified the rat model by integrating vascular endothelial growth factor receptor 2 (VEGFR2) inhibitor (SU5416) with a repeated injection of performed autologous clot segments (1-2 mm in length) weekly for 4 weeks. Rats were randomly assigned to Sham (saline injection), SU5416 (subcutaneous injection of SU5416 alone), Clots (w/SU) (Jugular vein injection of autologous clots generated from whole blood with SU5416), Clots + SU (Jugular vein injection of autologous clots combined with subcutaneous SU5416), and Clots (SU) + SU (Jugular vein injection of SU5416-containing autologous clots combined with subcutaneous SU5416) groups.

All rat procedures were performed under isoflurane anesthesia with 1.5 L/min oxygen flow. Once anesthesia was achieved, the clot segments (80-100) were aspirated into a 5-mL syringe with a 23 gauge needle. The clots in 2 mL of 10% tranexamic acid (TAX) solution, which suppresses fibrinolysis and stabilizes fibrin clots^3^, were then slowly injected into the right internal jugular vein. Simultaneously, SU5416 (20 mg/kg; MedChemExpress, Cat. #HY-10374, USA) in DMSO was administered via subcutaneous injection only during the first clot injection. The neck incision was sutured with an absorbable suture and disinfected with povidone-iodine. This procedure was repeated weekly for a total of four clot injections, while SU5416 was given only once. A detailed schematic of the procedure is shown in **Figure 2A**.

***Hemodynamic measurements*:** Two weeks after the final clot injection, a right heart catheterization procedure was performed to determine hemodynamic parameters. Under total anesthesia, abdominal cavity was exposed, and a catheter pre-filled with heparin was inserted into the right ventricle through the diaphragm using a 21G needle as previously described^4,5^. The right ventricular systolic pressure (RVSP) was measured using a pressure monitoring system and Labchart 8 software (AD Instruments, Sydney, Australia) at the Cardiovascular Research Core facility at the University of Kansas Meddical Center.

##### ***Echocardiographic assessment***:**** Transthoracic echocardiography was performed in rats with CTEPH and Sham controls using a high-resolution ultrasound system (Vevo F2, VisualSonics Inc.) equipped with a 46–57 MHz linear-array transducer. All measurements were obtained under light isoflurane anesthesia. Anesthesia was induced with 3% isoflurane and maintained at 1.0–1.5% to achieve a stable heart rate during imaging. Right ventricular wall thickness at end diastole (RVWTD) was measured from parasternal short-axis M-mode images. Pulmonary artery acceleration time (PAT) and pulmonary artery ejection time (PET) were obtained from pulsed-wave Doppler recordings of pulmonary artery flow, and the PAT/PET ratio was calculated as an index of pulmonary hemodynamics. Measurements were averaged from at least five consecutive cardiac cycles for each animal.

***Histological and immunohistochemical analysis*:** For immunofluorescent staining, the left lobe of lung tissue was perfused with 70% OCT in phostate-buffered saline (PBS) through tracheal catheterization and then embedded in OCT for cryosection. Lung sections (10 μm) were fixed with 4% paraformaldehyde, followed by blocking with 0.1% Triton X-100 and 5% normal goat serum at room temperature for one hour. After 3 washes with PBS, they were incubated with rabbit anti-human VWF immunoglobulin G (1:400) (Dako, Cat.#00073780, Glostrup, Denmark), anti-α-smooth muscle actin (α-SMA) (1:300) (Dako, Cat.#GA611,Glostrup, Denmark), anti-Fibrin (1:400) (Dako, Glostrup, Denmark), and anti-CD31 (1:200) (Santa Cruz Biotechnology, Cat.#sc-20071,TX, USA) at 4 °C overnight. After washing, the tissue sections were then incubated with a fluorescence-conjugated secondary antibody (FisherScientific, Carlsbad, California) at room temperature for one hour. Nuclei were counterstained with a mounting medium containing DAPI (Cell Signaling Technology, #8961S, MA, USA). Quantification of expression signals was performed blindly.

### For histology and other immunohistochemical studies, well-perfused lungs were fixed for 5 min by instillation of 10% PBS-buffered formalin through trachea catheterization at a transpulmonary pressure of 15 cm H_2_O, and then 48 h fixation at room temperature on a shaker. After paraffin processing, the tissues were cut into semi-thin 6 μm thick sections. Rat lung sections were then dewaxed and dehydrated. Antigen retrieval was performed by boiling the slides in an antigen retrieval buffer (Vector Lab, Newark, CA) for 1 minute in a high-pressure cooker. Thin sections were prepared using a microtome and stained with hematoxylin and eosin (H&E). Additionally, the adjacent sections were used for immunohistochemical staining for VWF (Dako, #00073780, Glostrup, Denmark), glycoprotein IIIa (Anti-β3/CD61, gift from Dr. Heyu Ni at University of Toronto,CA), fibrin (1:400) (Dako, Glostrup, Denmark), TGF-β1 (TGF beta 1 Antibody 3C11, Santa Cruz Biotechnology, Cat.#sc-130348, Dallas, TX, USA), PDGF-BB (PDGF-B Antibody, Santa Cruz Biotechnology, Cat.#sc-74494, Dallas, TX, USA), Endothelin-1 (Endothelin-1 Monoclonal Antibody, Invitrogen, Cat.#MA3-005, Thermo Fisher Scientific, MA, USA), and Ki-67 (Ki-67 Rabbit Monoclonal Antibody, clone D3B5, Cell Signaling Technology, Cat.#9129, MA, USA), followed by the peroxidase-conjugated secondary antibody and color reaction with 3,3′-diaminobenzidine/H2O2. Digital images were taken using a Carl Zeiss Axioplan light microscope (Göttingen, Germany).

### For assessment of vascular remodeling, the paraffin sections of lungs were dewaxed and dehydrated, and then stained with a Trichrome Masson Stain Kit (Sigma Aldrich, #HT15, MO, USA) as described previously. For assessment of pulmonary arterial (PA) wall thickness, PAs from images at 20x magnification were quantified by Image J. Wall thickness was calculated blindly as the distance between the internal wall and the external wall, divided by the distance between the external wall and the center of the lumen, as described previously. Measurements were conducted specifically in pulmonary arteries with a diameter of 50–100 µm.

***Real-time quantitative PCR*:** The livers of rats from different groups were collected and preserved in RNA later (Sigma Aldrich, #R0901, MO, USA). RNA was extracted and purified using PureLink RNA Mini Kit (Thermo Fisher Scientific, #12183018A, MA, USA) and reverse transcribed into cDNA using a SuperScript™ IV First-Strand Synthesis System (Invitrogen, CA, USA). Real-time quantitative PCR was performed using a QuantStudio 7 Flex Real-Time PCR System (Applied Biosystems). All primer sequences for RT-QPCR are detailed in the **Supplementary Table S1**.

***Single-cell RNA sequencing*:** Left lungs were harvested from two Sham rats and two CTEPH rats and immediately processed for single-cell RNA sequencing. Single-cell dissociation, library construction, and sequencing were performed by Singleron Biotechnologies (Connecticut, USA) using a standard 10x Genomics Chromium platform workflow. Briefly, viable single-cell suspensions from each sample were loaded onto the Chromium Single Cell Controller to generate barcoded single-cell libraries, which were subsequently sequenced on an Illumina platform with paired-end reads. Raw sequencing data were processed and analyzed using Seurat (v4) in R. Cells expressing fewer than 100 detected genes, cells with excessively high gene counts, or cells with a high proportion of mitochondrial transcripts were excluded from further analysis. After quality control, data from Sham and CTEPH groups were normalized and integrated using Seurat’s standard integration workflow. Principal component analysis (PCA) was performed for dimensionality reduction, and the integrated dataset was visualized using Uniform Manifold Approximation and Projection ^6^. Graph-based clustering was performed using the FindClusters function in Seurat. Marker genes used for lung cell type annotation are summarized in **Supplementary Table S2**. Marker genes used for pulmonary endothelial cell subtype annotation are summarized in **Supplementary Table S3**.

*Measurement of plasma biomarkers*: Plasma ADAMTS13 antigen levels in human samples were determined using a commercially available enzyme-linked immunosorbent assay (ELISA) kit (R & D Systems, Cat. #DADT130, Minneapolis, MN, USA), according to the manufacturer’s instructions. Human plasma samples, as described above, were thawed on ice and analyzed in duplicate. Absorbance was measured at 450 nm with wavelength correction at 570 nm using a microplate reader. ADAMTS13 antigen concentrations were calculated from a standard curve generated with recombinant human ADAMTS13 standards provided in the kit.

For biomarkers in rat plasma, blood was collected following the terminal procedure for RVSP measurement via the inferior vena cava. Whole blood was anticoagulated with EDTA and plasma was prepared by centrifugation at 3,000 rpm and stored at −80 °C until further analysis. Plasma levels of N-terminal pro–B-type natriuretic peptide (NT-proBNP) and D-dimer (D2D) were determined by the Rat NT-proBNP (Antibodies.com, #A74911, CA, USA) and Rat D2D ELISA Kit (Elabscience Bionovation, #E-EL-R0317, Hubei, China), respectively.

Plasma samples for TGF-β1 and PDGF-BB levels were obtained from rats of different experimental groups to avoid platelet platelet activation and artifactual release of platelet-derived growth factors. Briefly, following initial separation, plasma samples were further centrifuged at 10,000 × g for 10 min at 4 °C to remove any residual platelets and cellular debris. The resulting platelet-poor plasma was aliquoted and stored at −80 °C until analysis. Plasma levels of transforming growth factor–β1 (TGF-β1) were quantified using the Rat TGF-β1 Quantikine ELISA Kit (#DB100C) (R&D Systems, Minneapolis, MN, USA) according to the manufacturer’s instructions. Prior to measurement, plasma samples were subjected to acid activation to convert latent TGF-β1 into its immunoreactive form. Platelet-derived growth factor-BB (PDGF-BB) levels were determined using the Rat PDGF-BB Quantikine ELISA Kit (R&D Systems, Cat.#MBB00, Minneapolis, MN, USA) following the manufacturer’s protocol, without additional activation steps. All ELISA assays were performed and analyzed in a blinded fashion with respect to the experimental groups.

***In situ hybridization*:** *In situ* hybridization (ISH) was performed on paraffin-embedded liver and lung sections (10 µm) from Sham and CTEPH rats for detection of *Adamts13* and *vwf* mRNA expression. Tissue sections were deparaffinized in xylene, rehydrated through graded ethanol, and subjected to target retrieval and protease digestion following the manufacturer’s instructions (Advanced Cell Diagnostics, ACD Bio, USA). ISH was carried out using RNAscope probes specific for rat *Adamts13* (RNAscope *Adamts13* probe (1802671-C1) (ACD Bio USA, Newark, CA) and rat *Vwf* (RNAscope *Vwf* probe, #413401) (ACD Bio, USA). Probes were hybridized at 40°C for 2 h, followed by sequential amplification steps using the RNAscope detection system. Signals were visualized using fluorescent labeling methods. A positive control probe (PPIB) and a negative control probe (dapB) were included in each run to ensure assay quality. Sections were counterstained with DAPI, mounted, and imaged using bright-field or fluorescence microscopy. For quantification, ISH signals were analyzed in a blinded manner using ImageJ and expressed as dots per cell or percentage of positive cells.

*Assessment of plasma ADAMTS13 Activity*: The proteolytic activity of recombinant ADAMTS13 in rat plasma was determined using the FRETS-VWF73 substrate as previously described^7,8^. Rat plasma endogenous ADAMTS13 activity was measured using the Cattle FRETS-VWF71 substrate (kindly provided by Dr. Joshua Muia at Versiti, Milwaukee, WI)^9^. A pooled plasma sample prepared from several WT rats served as the calibration standard, defined 100% activity.

##### ***Flow cytometric analysis*:** Whole blood was collected from rats via retro-orbital sinus bleeding into tubes containing acid–citrate–dextrose (ACD) solution B (#786-494) (G-Bioscience, St. Louis, MO) as an anticoagulant. Platelet-rich plasma (PRP) was obtained by centrifugation at 100 × g for 10 min at room temperature without brake. PRP was further centrifuged at 800 × g for 10 min to pellet platelets. The platelet pellet was gently resuspended in a calcium-free Tyrode’s buffer supplemented with prostaglandin E1 (1 μmol/L). Platelets were then stained with a FITC-conjugated anti-CD61 antibody to identify platelets (#104306-BL) (BioLegend, San Diego, CA) and a PE-conjugated anti-P-selectin(CD-62P) antibody to assess platelet activation (#10019-154) (BioLegend, San Diego, CA) for 20 min in the dark at room temperature. As described above, unstained platelets, as well as platelets stained with either anti-CD61 or anti-CD62P antibody alone, were used as controls. Samples were diluted with Tyrode’s buffer and analyzed immediately using a spectral flow cytometer (Cytek Aurora, Cytek Biosciences, Fremont, CA). Platelets were gated based on forward scatter (FSC) and side scatter (SSC) characteristics and further identified by CD61 positivity. At least 10,000 CD61-positive platelet events were acquired per sample. Data were analyzed using FlowJo software (version 10.8.1). Platelet activation was expressed as the percentage of CD62P-positive platelets.

*Microfluidic shear-based assay*: Microchannels (Fluxion Bioscience, San Francisco, CA) were coated with a type 1 fibrillar collagen (100 μg/mL) (Chrono-Log, Overland Park, KS) in 0.01M HCl as previously described^10^. The surface was blocked with 0.5% bovine serum albumin in PBS. Whole blood collected from rats was anticoagulated with a thrombin inhibitor, D-phenylalanyl-L-prolyl-L-arginine chloromethyl ketone (PPACK) (100 μM) (Sigma-Aldrich, St. Louis, MO) and perfused under arterial shear (50 dyne/cm^2^) over collagen-coated surfaces at the same time. The thrombus formation was recorded every 3 seconds for 180 seconds. The rate and total surface area coverage of fluorescent platelets were determined using the Montage software (Fluxion Bioscience, San Francisco, CA) off-line. Prior to the initiation of perfusion, a PE-conjugated anti-CD61 antibodies (#104308-BL) (BioLegend, San Diego, CA) or a FITC-conjugated-anti-CD61 (#104306-BL) (BioLegend, San Diego, CA) was added to the blood samples 10 minutes before the experiment to label circulating platelets. At the end of the perfusion, microchannels were gently rinsed with 4% PFA to fix adherent cells and thrombi, followed by blocking with a BSA-containing buffer to reduce nonspecific binding. Subsequently, channels were incubated with additional fluorescently conjugated antibodies, including Alexa Fluor® 488–conjugated anti–VWF antibody (#EPR25069-131) (Abcam, Cambridge, UK) and APC-conjugated anti–P-selectin (#10019-152) (Invitrogen, Carlsbad, CA) as needed. Images were then acquired using a confocal laser scanning microscope under identical settings across samples.

***VWF multimers assay*:** Plasma VWF multimers were analyzed by Western blotting following electrophoresis on a 1% agarose gel and being transferred to a nitrocellulose membranes (Bio-Rad, CA). VWF expression was detected by immunoblotting with rabbit anti-human VWF immunoglobulin G (Dako, Glostrup, Denmark) as previously described^11^.

*SDS-PAGE and Western blotting*: Western blotting was performed using rabbit anti-human VWF immunoglobulin G (Dako, Cat.#00073780, Glostrup, Denmark), anti-P-selectin (Ebioscience, San Diego, CA), anti–TGF-β1 (Santa Cruz Biotechnology, Cat.#sc-130348, Dallas, TX), and anti–PDGF-BB (Santa Cruz Biotechnology, Cat.#sc-74494). The β-actin as the internal loading control for all samples was detected with anti- β-actin IgG (Cell Signaling Technology, Cat.#3700S, Danvers, MA). HRP-conjugated Goat-Anti-Rabbit IgG (Cell Signaling Technology, Cat.#7074) served as the secondary antibody. Protein lysates from lung tissues were prepared by mincing samples in cold RIPA lysis buffer (MilliporeSigma, Cat.#20-188, St. Louis, MO) supplemented with protease inhibitor cocktails (Roche, Cat.#04693132001, Basel, Switzerland). The lysates were thoroughly homogenized and subjected to ultrasonic disruption. For washed rat platelets, cells were directly lysed in cold RIPA buffer supplemented with protease inhibitors and incubated on ice for 10 minutes without homogenization or sonication. Subsequently, NuPAGE LDS Sample Buffer (Invitrogen, Cat.#NP0007, Carlsbad, CA) was added to all samples, followed by heating at 70 °C for 20 minutes prior to electrophoresis. Equal amount of protein was loaded for 10% SDS-PAGE and Western Blotting. Transfer using PVDF membranes (ThermoFisher Scientific, Cat.#88518, Waltham, MA). Detection was carried out with the Odyssey imaging system (LI-COR Biosciences, Lincoln, NE). The quantification of densitometric measurements for blot images was performed on the Image J.

##### *Administration of human recombinant ADAMTS13 in rats*: Recombinant ADAMTS13 variant truncated after the 5th TSP1 repeat (rA13-T5) that is proteolytically active in cleaving VWF was used for therapeutic intervention in CTEPH rats^12^. A schematic comparison of the molecular structures is provided in Supplementary Figure S6A. Prior to in vivo administration, the amount and structural integrity of rA13-T5 were confirmed by Western blotting (Supplementary Figure S6B) and its proteolytic activity by FRETS-VWF73 assay.

Rats were randomly assigned to three experimental groups: 1. Sham-operated rats receiving vehicle (PBS) treatment (Sham group); 2. rats with CTEPH receiving vehicle treatment (CTEPH group); and 3) rats with CTEPH treated with rA13-T5 (rA13-treated group). A vehicle or rA13-T5 was administered intravenously at 40 IU/kg, a dose used for treatment of patients with congenital TTP^13^, every week for 4 weeks, similar to the schedule for induction of CTEPH in rats.

*Tail bleeding time assay*: Tail bleeding time was assessed as an in vivo measure of primary hemostasis. Rats were restrained and anesthetized with inhaled isoflurane. After disinfection with 70% ethanol, a 3-mm segment of the distal tail was amputated using a sterile blade. Timing began immediately, and bleeding was monitored continuously. Blood was gently blotted with filter paper without contacting the wound. Bleeding time was defined as the interval from transection to complete cessation of visible bleeding, with bleeding stopped by gentle pressure if a preset cutoff time was exceeded. Measurements were performed after the final rADAMTS13 administration.

*Statistical analysis*: Statistical analysis was performed on Prism 9 (Graphpad, Bonston, MA). All data are presented as individual values ^14^, means ± standard errors of the means (SEM). P values were calculated using two-tailed Student’s t test for data that passed the normality test or the Mann-Whitney rank-sum U test for data that were not normally distributed for comparison between two groups. One-way analysis of variance (ANOVA) with Tukey’s test was used to evaluate differences among three or more groups. Two-way mixed-effects ANOVA followed by Tukey’s test was used for experiments involving repeated measurements and multiple groups. A P-value less than 0.05 or less than 0.01 was considered statistically significant or highly significant.

**Supplementary Tables**

**Table S1. Primer sequences used for quantitative PCR and genotyping**

| **For Q-PCR**: | | |
| --- | --- | --- |
| *Vwf* | Forward primer: 5'-GAG CAG ATC CAT CCA GCA TT-3'  Reverse primer: 5'-GTG AGG GCC AGA ACT AAC AG-3' | Integrated DNA technologies |
| *Beta-actin* | Forward primer: 5'-CGC AAG TAC TCT GTG TGG AT-3'  Reverse primer: 5'-GTA AAA CGC AGC TCA GTA ACA G-3' | Integrated DNA technologies |
| *Adamts13* | Forward primer: 5'-GTG CAA GAG TGT TTT GGA GC-3'  Reverse primer: 5'-GTT TTG AGA CGA AAC GCC TG-3' | Integrated DNA technologies |
| **For genotyping**: | | |
| *Adamts13* | Forward primer: 5'-ATG GGC ACT CTC TTC ACC AC-3'  Reverse primer: 5'-CAG CTT CCC CTA CCA AGA CA-3' | Integrated DNA technologies |
| *Vwf* | 3’ End PCR (250 bp)  VWF WT F: cggcagactcctactgctac  VWF 3G R: attagatggcctcagcctcca  Deletion PCR (330 bp)  VWF 5G F:   tggactggactgtcatgggaaa  VWF 3G R:  attagatggcctcagcctcca  5’ End PCR (350 bp)  VWF  5G F:  tggactggactgtcatgggaaa  VWF WT R:  ctggatggatctgctcaggc | Integrated DNA technologies |

**Table S2. Marker genes used for lung cell type annotation in**

**single-cell RNA sequencing analysis**

| Cell type | Marker genes |
| --- | --- |
| Alveolar macrophages | Adgre1 (F4/80), Lyz2, Cd68, Csf1r |
| Alveolar type I epithelial cells (AT1) | Ager, Hopx, Pdpn |
| Alveolar type II epithelial cells (AT2) | Sftpc, Sftpb, Sftpa1, Sftpd |
| Fibroblasts | Col1a1, Col1a2, Dcn, Lum |
| Smooth muscle cells (SMCs) | Acta2, Tagln, Myh11 |
| Endothelial cells (ECs) | Pecam1, Cdh5, Kdr |
| Neutrophils | S100a8, S100a9 |
| Mononuclear phagocytes | Lyz2, Fcgr3, Ctss |
| Erythrocytes | Hbb-bs, Hba-a1 |
| Erythroid progenitor cells | Gata1, Klf1 |
| T cells | Cd3d, Cd3e, Trbc1 |
| B cells | Cd79a, Ms4a1 |
| Ciliated cells | Foxj1, Tekt1 |
| Club cells | Scgb1a1, Scgb3a1 |
| Proliferating club cells | Scgb1a1, Mki67, Top2a |

##### Table S3. Marker genes for pulmonary endothelial cell subpopulations

| Endothelial cell subtype | Marker genes |
| --- | --- |
| Arterial endothelial cells | Gja5, Gja4, Hey1, Sema3g, Igfbp3 |
| Capillary endothelial cells | Ednrb, Rgcc, Plvap, Ramp3, Gpihbp1, Kdr, Cxcl12, Hpgd |
| Venous endothelial cells | Ackr1, Selp, Icam1, Nr2f2, Vwf, Nrp2, Vcam1, Cpe, Mmrn1, Nrp1 |
| Lymphatic endothelial cells | Pdpn, Prox1, Reln, Ccl21 |

**Table S4 Demographic and clinical information for all participants**

| CTEPH patients | |  |  |  |  |  |
| --- | --- | --- | --- | --- | --- | --- |
| PHBR # | Diagnosis | Age | Gender | Race | Ethnicity | History of Treatment Prior to Lab draw |
| 010 | CTEPH | 64 | M | White | Non Hispanic | Adempas 7.5 mg |
| 034 | CTEPH | 64 | M | White | Non Hispanic | Adempas 7.5 mg |
| 114 | CTEPH | 48 | F | White | Non Hispanic | Adempas 7.5 mg Opsumit 10 mg |
| 178 | CTEPH | 26 | F | White | Non Hispanic | Letairis 10 mg |
| 050 | CTEPH | 62 | F | White | Non Hispanic | Adempas 7.5 mg Remodlin IV 2 ng/kg/min |
| 097 | CTEPH | 57 | F | White | Non Hispanic | Adempas 7.5 mg |
| 138 | CTEPH | 71 | M | White | Non Hispanic | None |
| 140 | CTEPH | 60 | M | White | Non Hispanic | None |
| 158 | CTEPH | 58 | F | White | Non Hispanic | Adempas 7.5 mg |
| 179 | CTEPH | 62 | M | White | Non Hispanic | Adcirca 40 mg Letaris 10 mg Inhaled Tyvaso 256 mcg |
| 214 | CTEPH | 57 | M | White | Non Hispanic | Adempas 1 mg |
| 229 | CTEPH | 57 | M | White | Non Hispanic | Revatio 60 mg |
| 20610 | CTEPH | 58 | Female | White | Not Hispanic or Latino |  |
| 15267 | CTEPH | 62 | Female | White | Not Hispanic or Latino |  |
| 10644 | CTEPH | 76 | Female | White | Not Hispanic or Latino |  |
| 2977 | CTEPH | 63 | Female | White | Not Hispanic or Latino |  |
| 14219 | CTEPH | 77 | Male | White | Not Hispanic or Latino | Adempas 7.5 mg |
| 29283 | CTEPH | 65 | Female | White | Not Hispanic or Latino |  |
| 36022 | CTEPH | 65 | Female | White | Not Hispanic or Latino |  |
| 27462 | CTEPH | 38 | Female | White | Not Hispanic or Latino | Adempas 7.5 mg Opsumit 10 mg |
| IPAH patients | |  |  |  |  |  |
| 003 | Idiopathic PAH | 33 | F | White | Non Hispanic | Remodulin IV 35ng/kg/min Macitentan 10mg |
| 004 | Idiopathic PAH | 41 | F | White | Non Hispanic | Revatio - 20mg TID Remoduin IV 22ng/kg/min |
| 008 | Idiopathic PAH | 60 | F | White | Non Hispanic | Tyvaso Nebulizer 1.74 mg/2.9 mL 12puffs QID Revatio 20mg TID Letairis 10mg |
| 009 | Idiopathic PAH | 80 | M | White | Non Hispanic | Tyvaso Nebulizer 1.74 mg/2.9 mL 12puffs QID |
| 020 | Idiopathic PAH | 48 | F | White | Non Hispanic | Adcirca 40mg Letairis 10mg Sotatecept 0.7mg/kg |
| 023 | Idiopathic PAH | 31 | F | White | Non Hispanic | Revatio - 20mg TID Remodulin SQ 34.9 ng/kg/min |
| 024 | Idiopathic PAH | 62 | F | White | Non Hispanic | Tyvaso Nebulizer 9 breaths QID Revatio 20mg TID |
| 048 | Idiopathic PAH | 35 | F | White | Non Hispanic | Adcirca 40mg Veletri 28ng/kg/min |
| 182 | Idiopathic PAH | 57 | F | White | Non Hispanic | Orenitram 0.625mg |
| Healthy Donors | |  |  |  |  |  |
|  | Diagnosis | Age | Gender | Race | Ethnicity |  |
| #1 | Donor | 72 | Male | White | Not Hispanic or Latino |  |
| #2 | Donor | 68 | Female | White | Hispanic or Latino |  |
| #3 | Donor | 32 | Female | White | Not Hispanic or Latino |  |
| #4 | Donor | 63 | Female | White | Not Hispanic or Latino |  |
| #5 | Donor | 59 | Female | White | Not Hispanic or Latino |  |
| #6 | Donor | 62 | Female | White | Not Hispanic or Latino |  |
| #7 | Donor | 76 | Female | White | Not Hispanic or Latino |  |
| #8 | Donor | 63 | Female | White | Not Hispanic or Latino |  |
| #9 | Donor | 77 | Male | White | Not Hispanic or Latino |  |
| #10 | Donor | 67 | Female | White | Not Hispanic or Latino |  |
